# Root Age Structure Governs Long-Term Architecture and Hydraulic Function in Perennial Plants: a Modelling Framework

**DOI:** 10.64898/2026.10.03.756415

**Authors:** Lukas Fichtl, Mona Giraud, Daniel Leitner, Andrea Schnepf, Katharina Steng, Matthias Friedel

**Author notes:** These authors contributed equally to this work (Lukas Fichtl, Mona Giraud).

## Abstract

**Background and aims:** Perennial root systems develop over decades and exhibit continuous turnover of structural and absorptive components, yet most root architecture models treat woody roots as permanent and neglect root age structure. We developed an empirically constrained modelling framework that explicitly represents age-structured demography, growth regulation and dormancy in perennial root systems.

**Methods:** We extended the functional–structural plant model CPlantBox to simulate perennial root development over decades by integrating age-dependent survival, secondary radial growth and climate-driven dormancy. The model was parameterised using unique three-dimensional root system data from young and mature grapevines, together with dendrochronological root age measurements, and coupled to root hydraulic properties.

**Key results:** Empirical analyses revealed multi-cohort age distributions and substantial turnover of lignified structural roots, contradicting the assumption of a static woody scaffold. The model reproduced age distributions, order-specific survival probabilities and fine-to-structural root length ratios, and generated realistic, genotype-specific three-dimensional architectures. Simulated uptake profiles and hydraulic scenarios showed strong age- and architecture-driven effects.

**Conclusions:** Explicitly representing woody root age distributions and age-dependent processes may fundamentally change predictions of perennial root system architecture and function. The framework advances root modelling beyond annual paradigms and provides a transferable tool for analysing long-term architectural and hydraulic consequences of genotype × climate interactions in perennial crops.

## 1 Introduction

Perennial plants dominate most terrestrial ecosystems and contribute substantially to global agricultural production (Poppenwimer et al. 2023; Scott et al. 2022), yet the long-term development of their root systems remains poorly understood (Freschet et al. 2021). Unlike annuals, perennial root systems persist for decades and are shaped by the continuous emergence, maturation and loss of both absorptive and woody components (Munné-Bosch 2014; Roumet et al. 2006). These dynamics imply that perennial root system architecture (RSA) is inherently demographic rather than purely structural.

RSA governs where and how effectively plants acquire water and nutrients, explore soil volumes, store carbon and ensure mechanical stability (de Dorlodot et al. 2007; Lynch 1995). Mechanistic root models and functional–structural plant models (FSPMs) enable explicit simulation of growth and plant–soil interactions in space and time (Diggle 1988; Leitner et al. 2010; Lynch et al. 1997), and three-dimensional approaches have become increasingly important for crop improvement and ecosystem prediction (Jourdan and Rey 1997; Morris et al. 2017). However, most existing RSA models were developed for annual species and focus on a single growing season (Dupuy et al. 2010), leaving perennial root systems poorly parameterised and mechanistically underrepresented (Barczi et al. 2018; Cochard et al. 2021; Fichtl et al. 2023; Saint Cast et al. 2019a).

A central limitation is the lack of explicit representation of root age distributions, age-dependent growth potential, seasonal dormancy and long-term (decadal) turnover, preventing models from reproducing the demographic steady states observed in mature perennials (Freschet et al. 2021). Developing perennial frameworks is challenging because empirical constraints on age-dependent growth, mortality, secondary thickening and hydraulic transitions remain scarce (Pierret et al. 2007), and mature root systems are difficult to access and rarely excavated at sufficient depth or replication (Saint Cast et al. 2019b).

Perennial plants differ fundamentally from annuals in their architectural and functional dynamics. Structural roots undergo secondary thickening, lignification and hydraulic maturation (Gambetta et al. 2013; Strock and Lynch 2020), while absorptive fine roots turn over rapidly and respond to environmental variation (Anderson et al. 2003; Sun et al. 2016). Whether lignified structural roots form a quasi-permanent scaffold or are continuously renewed over decades remains unresolved (Munné-Bosch 2014), with important implications for hydraulic functioning, carbon residence and drought resilience. In addition, perennial growth is shaped by seasonal dormancy and episodic growth pulses (McCormack et al. 2015), processes not captured by simply extending annual models in time.

Although several models have been extended to simulate perennial plants, perenniality is typically represented through root-type classifications or predefined lifespan rules rather than emerging from empirically constrained, age-structured demography (Collet et al. 2006; Jourdan and Rey 1997; Saint Cast et al. 2019b; Vercambre et al. 2003). Consequently, structural turnover, age-dependent growth regulation, dormancy and their hydraulic consequences are rarely integrated within a unified framework.

Addressing this gap requires modelling approaches that place root age distribution at the centre of system dynamics, explicitly linking age-dependent growth, secondary development, dormancy and hydraulic function. Perennial RSA typically progresses from an exploratory establishment phase to a stabilised framework dominated by fine-root turnover and localised remodelling, reinforcing the view of perennial root systems as age-structured populations rather than static architectures (Bernardo et al. 2025).

Here, we develop a perennial root system modelling framework that integrates age-structured demography, three-dimensional architecture and hydraulic function. Using grapevine (*Vitis vinifera* L.) as a model woody perennial, we combine empirical datasets from young (1–2 years) and mature (48 years) field-grown plants with dendrochronological root age reconstructions to parameterise and evaluate long-term survival, growth regulation and secondary development. Grapevine provides a suitable system because belowground performance is strongly shaped by rootstock-specific RSA and hydraulic traits (Bernardo et al. 2025; Delrot et al. 2020; Keller 2010).

Our objectives were to: (i) establish root age structure as an organising principle of perennial RSA; (ii) develop a transferable, process-based framework for multi-decadal root system simulations; and (iii) assess how the resulting age-dependent architecture of the vine root system shape water uptake patterns and hydraulic responses under contrasting climatic scenarios. By linking empirical root demography to architecture and function, this study advances perennial root modelling beyond static lifespan assumptions and provides new insights into the long-term functioning of woody root systems.

## 2 Materials and Methods

The Materials and Methods comprised three components: (i) acquisition and analysis of empirical root system architecture (RSA) and root age data (section 2.1), (ii) derivation of demographic and growth parameters (section 2.2), and (iii) implementation and calibration of the perennial modelling framework (section 2.3).

### 2.1 Experimental Data

Root system architecture of young and mature ‘Riesling’ grapevines was characterised by destructive excavation and three-dimensional (3D) digitisation under field conditions. Root diameter measurements and dendrochronological cross-sections were used to estimate root age and secondary growth patterns.

#### 2.1.1 Young grapevine root system architecture

Young-vine RSA data originated from a vineyard establishment experiment at Hochschule Geisenheim University, Germany, described in detail in Fichtl et al. (2025). Following standard bench grafting procedures (Waite et al. 2015), cuttings for the scion (upper part of the grafting) of the new graftings were obtained from “Riesling” vines (Vitis vinifera L.; clone N90). Cuttings from three contrasting grapevine varieties (see below) were used for rootstock (lower part of the grafting). The cuttings were grafted together and grown for one year in a nursery. Following this, the adventitious roots were cut at 10-15cm of length and then planted in the field, standardising the resource available to the plant at the start of the experiment (Bates et al. 2002). Vines were planted in a randomised complete block design and excavated at 3, 6, 15 and 18 months after planting (n = 96; 24 per time point). Root systems were manually excavated and digitised in 3D using an electromagnetic tracking system (Polhemus Fastrak, Colchester, USA) following Fichtl et al. (2024). Root length was calculated from the digitised geometries. The dataset is publicly available (Fichtl and Friedel 2025b).

The three rootstock varieties were ‘101-14’, ‘SO4’, and ‘R110’ (parentage in Fichtl et al. (2025)). Those three genotypes have different level of resistance to drought (Keller 2010): R110 is rather resistant while SO4 is strongly sensitive to drought, while 101-14 is somewhat sensitive to drought stress.

#### 2.1.2 Mature grapevine root system architecture

Although no mature rootstocks of the same genotypes with an age sufficiently advanced were available, an example dataset for older root systems was produced using three mature (48-year-old) vines. ‘Riesling’ grapevines grafted onto rootstock ‘5C’ were excavated in Oestrich-Winkel, Germany (Friedel et al. 2020). The extraction was conducted in winter (outside of the vines’ growing season). Following our assumptions regarding the life-cycle of unsuberised roots (see Section 2.3.3), these roots (thin roots and young structural roots) died before the excavation. Therefore, although it was not observed on the excavated root systems, we assumed they were present during the previous growing season. Moreover, excavations were spatially restricted (1.60 m × 2.00 m; 1.50 m depth), and roots extending beyond this volume were truncated at the excavation boundary. Exposed root systems were digitised using the same 3D acquisition protocol as for young vines (Fichtl et al. 2024), and root lengths were derived from the digitised geometries. The dataset is publicly available (Fichtl and Friedel 2025a). In terms of genotypes, ‘5C’ is most similar to ‘SO4’, and they both share a high sensitivity to drought. (Keller 2010).

#### 2.1.3 Root age estimation

All trunk-borne (“main”) roots of mature vines were sampled for age determination. Basal and distal diameters were measured; for truncated roots, distal measurements were taken at the excavation cut surface. For lateral roots emerging from these axes, basal diameter only was recorded. To account for irregular cross-sections, diameter was calculated as the mean of maximum and minimum measurements. These measurements formed the basis for dendrochronological age estimation, age–diameter relationships, and subsequent analyses of secondary growth and structural root age distributions.

### 2.2 Root Demography and Process Parameters

Woody root age was determined dendrochronologically. After 3D digitisation, roots were sampled at basal and distal positions, with tissue sections of approximately 5 cm length taken at each position, soaked in water for five days, sectioned manually, and examined under a binocular microscope (ZOOM 2000, Leica Microsystems, Germany). Annual growth rings were clearly visible under transmitted light without staining and were counted directly. Ring counts were related to measured diameters to derive age–diameter relationships. To validate ring-based age estimates, additional roots of known age (1–4 years) from young vines were analysed analogously.

Based on the dendrochronological dataset from mature vines, four key parameter groups were derived for the perennial model: (i) structural root age distributions; (ii) age-dependent survival and turnover functions; (iii) age-dependent damping of elongation (Relative Growth Potential, RGP); and (iv) age-dependent secondary radial growth.

#### 2.2.1 Structural root age distributions

Structural root age distributions were derived from dendrochronological age estimates of trunk-borne roots. Age frequencies were first summarised per vine and then pooled across individuals to obtain overall distributions. Linear age–diameter relationships were fitted by regression and used to parameterise survival, turnover and age-dependent growth processes.

#### 2.2.2 Survival and structural root turnover estimation

Structural root survival was analysed separately for “main”, first-order lateral (“sub”), and higher-order lateral (“subsub”) roots. Ages of “main” roots were determined directly from annual ring counts, whereas ages of “sub” and “subsub” roots were estimated using the fitted age–diameter relationship.

Empirical survival curves were obtained using nonparametric Kaplan–Meier estimators (Kaplan and Meier 1958). Parametric Weibull models (Weibull 1939) were fitted to derive continuous, order-specific survival functions of the form

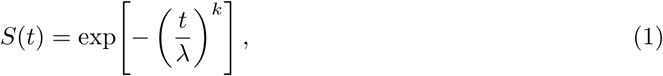

where *λ* (scale parameter in BEDD) and *k* (shape parameter, dimensionless) were estimated from the empirical age data (see Section 3.2.2). Parameters were derived from “main” roots and applied to first- and higher-order laterals.

#### 2.2.3 Relative Growth Potential as an age-dependent damping function

We assumed that all roots followed an asymptotic growth function of the form:

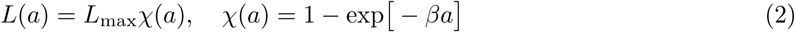

where *a* denotes root age (years), *β* (*year^−^*^1^) an initial growth parameter, *L*(*a*) (*cm*) the root length at age *a*, and *L*_max_ (*cm*) the maximum attainable root length. To quantify age-dependent decline in root elongation, the function *χ*(*a*) introduced a Relative Growth Potential (RGP) describing the reduction in axial growth capacity with increasing root age. The RGP parameters were derived from combined dendrochronological age estimates and 3D architectural measurements of excavated “main” roots.

Basal and distal segment ages (*a_b_* and *a_d_*) were determined from ring counts and age–diameter relationships, respectively, and related to the corresponding 3D segment lengths. Root length *L* (*cm*) was modelled as the increment of a cumulative age-dependent growth function,

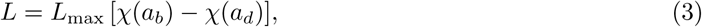

Parameters were estimated using nonlinear least squares.

In a subsequent step (section 2.3.9), age-dependent growth was mapped onto biologically effective degree days (BEDD, see Supplementary Methods S2) to obtain climate-driven elongation functions for dynamic simulations.

#### 2.2.4 Age-dependent secondary radial growth

Secondary radial growth was quantified by relating basal root diameter to ring-counted root age. For structural roots, segment diameter *D* (*mm*) was modelled as a linear function of its age *a*_seg_, (*year*),

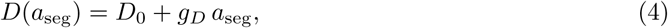

where *D*_0_ (*mm*) is the intercept and *g_D_* (*mm year^−^*^1^) the annual radial increment. The slope *g_D_* represents the empirical rate of secondary thickening and was used to parameterise age-dependent radial growth of structural roots in the perennial model (see Supplementary Methods S4).

### 2.3 Perennial Model Extensions

Figure 1 summarises the key perennial extensions implemented in CPlantBox. In the subsequent subsections, each of the components is described in detail, including its empirical foundation, mathematical formulation, and integration into the perennial CPlantBox framework.

**Figure 1:**
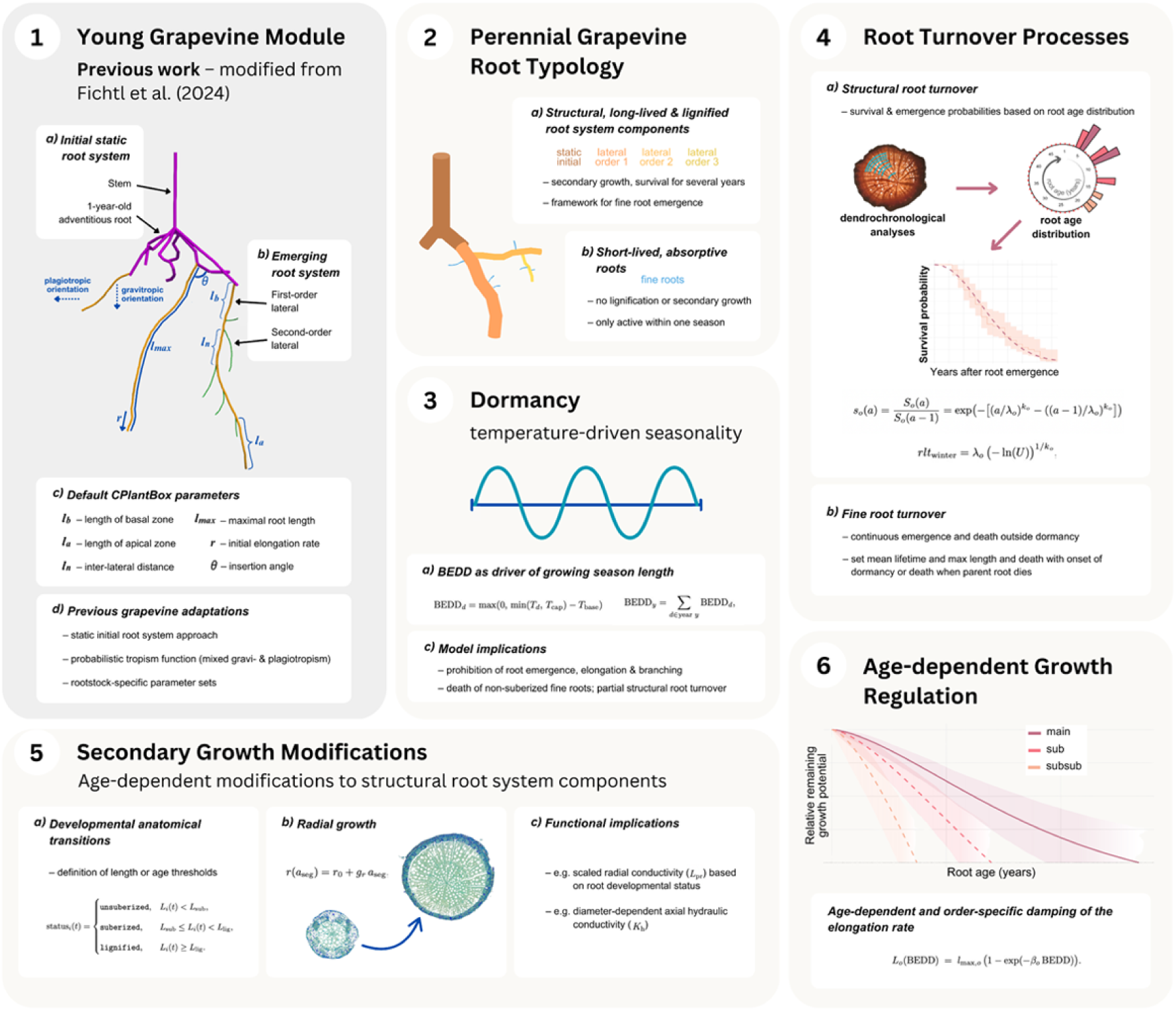
Conceptual overview of the perennial extensions implemented in CPlantBox. The framework integrates (1) prior grapevine-specific adaptations (section 2.3.1); (2) a perennial root system typology distinguishing structural and absorptive components (section 2.3.2); (3) a temperature-driven dormancy formulation (Section 2.3.3); (4) age-dependent structural and fine-root turnover processes (section 2.3.4 and section 2.3.5); (5) secondary growth and developmental status transitions (section 2.2.4, section 2.3.6, and section 2.3.7); and (6) age-dependent growth regulation (section 2.3.8 and section 2.3.9).

#### 2.3.1 Model framework and prior grapevine-specific adaptations

Perennial extensions were implemented within the functional–structural modelling framework CPlantBox (Giraud et al. 2023; Zhou et al. 2020), which represents plants as three-dimensional networks of connected segments following the CRootBox formalism (Schnepf et al. 2018). The framework supports rule-based growth, branching, secondary development and organ turnover, and can be coupled to hydraulic modules (Giraud et al. 2023).

CPlantBox was previously adapted to simulate grapevine root system establishment across root-stocks (Fichtl et al. 2024), including initialisation from empirically measured one-year-old root systems and probabilistic tropism functions reproducing genotype-specific growth orientations. The present study builds on this baseline and extends it to fully age-structured, multi-decadal perennial simulations.

#### 2.3.2 Root typology and branching rules

Following the convention of FSPMs (Dunbabin et al. 2013; Morandage et al. 2022; Schnepf et al. 2018; Zhou et al. 2020), we implemented a developmental classification of root typology, distinguishing (i) a persistent perennial framework (stem and trunk-borne axes; orders 0–1), (ii) long-lived structural roots (orders 2–6), and (iii) short-lived absorptive fine roots (orders *≥*7; Figure 2). Structural roots undergo maturation and secondary thickening and are subject to age-dependent survival and growth regulation, whereas fine roots are non-lignifying terminal modules with rapid turnover. We capped the maximum order of structural roots at 6 for two reasons: we did not observe higher structural root order in our experimental dataset, and when higher orders were permitted, structural roots of order *>* 6 essentially failed to emerge over the 50-year simulation.

**Figure 2:**
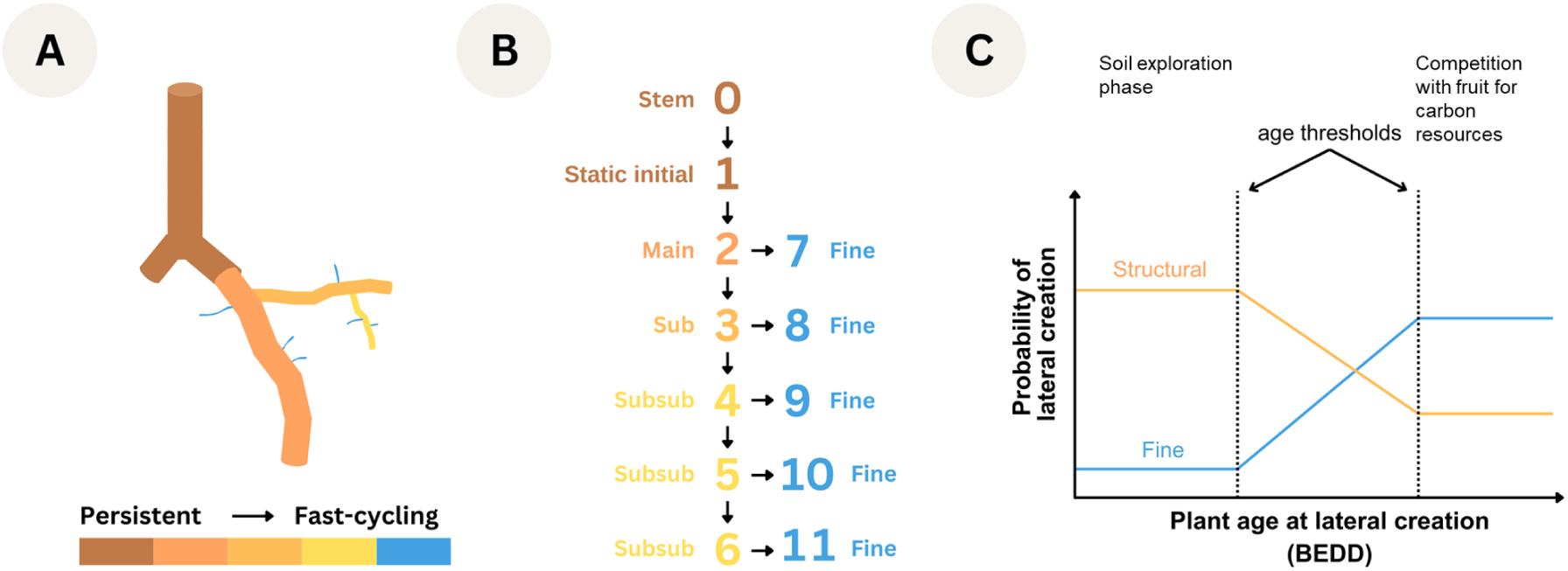
Implementation of root typology and branching rules in the perennial model. (A) Schematic representation of persistent structural roots and fast-cycling fine roots (color gradient indicates the transition from persistent to fast-cycling). (B) Branching rules between root orders, showing the order-specific identity of potential laterals (structural vs. fine). (C) Assumptions regarding the age-dependent branching probabilities for lateral creation: the probability of producing structural versus fine laterals changes with plant age at the time of lateral initiation (BEDD) according to specified age thresholds.

Order 1 represents trunk-borne axes present at planting. As these roots were cut, they do not grow during the experiment, but their secondary growth and death is simulated. Structural laterals can grow out of these static roots but no fine roots. Order 2 includes new trunk-borne roots and primary laterals (“main”), order 3 major laterals (“sub”), and orders 4–6 distal structural laterals (“subsub”). Fine roots (orders 7–11) emerge exclusively from living structural segments and do not branch further.

Branching is governed by CPlantBox successor tables specifying allowed successor types, maximum lateral numbers per segment, and age-dependent probabilities of initiating structural versus fine-root laterals (Supplementary Methods S1). Successor relations were restricted to biologically plausible transitions, i.e., structural roots can have both structural and fine roots as laterals, while fine roots can only have other fine roots as laterals. Age-dependent branching probabilities were implemented to represent developmental shifts from early exploratory structural expansion to increasing fine-root allocation and eventual architectural stabilisation.

Together, this typology and branching scheme provide a mechanistic representation of perennial root demography, allowing structural persistence and fine-roots’ turnover to emerge from age-dependent local rules.

#### 2.3.3 Dormancy

Root growth was restricted to an annually recurring active season, consistent with studies in temperate vineyards reporting pronounced spring and post-harvest fine-root flushes and negligible winter activity (Bauerle et al. 2008; Comas et al. 2010). Limited winter root growth has been observed in warmer climates (Mahmud et al. 2023), but was not considered under the cool-temperate conditions of this study.

Dormancy was implemented using a temperature-driven formulation based on biologically effective degree days (BEDD; Gladstones (2021); Supplementary Methods S2). Annual heat sums define the active growth period during which elongation, branching and fine-root production are permitted; outside this window, growth is suppressed.

At dormancy onset, newly formed unsuberised fine roots are removed (i.e., roots belonging to the fine root class and roots belonging to the structural root class but not yet suberised), consistent with the absence of white roots in winter excavations. Structural roots persist across dormancy according to age-dependent survival. This formulation links seasonal root activity directly to climatic forcing without relying on fixed calendar dates.

#### 2.3.4 Structural root turnover, emergence and secondary growth

Turnover of structural roots was implemented as a stochastic, age-dependent survival process. Upon creation, each structural root is assigned an individual lifetime drawn from an order-specific Weibull distribution parameterised from dendrochronological age data (Supplementary Methods S3). Roots are removed, together with their downstream subtree, once their age exceeds the assigned lifetime. In addition, newly formed roots that fail to reach a minimum length before dormancy are removed to ensure winter persistence of sufficiently developed structural axes.

Structural root emergence is governed exclusively by the successor-based branching rules described above (section 2.3.2; Supplementary Methods S1). New structural roots arise probabilistically from existing structural segments according to age-dependent branching probabilities, without explicit demographic balancing. Consequently, population-level age structures emerge from the interaction between stochastic survival and local branching dynamics.

Directional growth follows a probabilistic tropism function combining gravitropic and plagiotropic tendencies, as previously implemented for grapevine in CPlantBox (Fichtl et al. 2024). Order- and genotype-specific weighting reproduce the coexistence of vertically oriented and laterally spreading structural roots observed in the empirical dataset.

Secondary radial growth of structural roots was implemented as an age-dependent linear function at the root segment level, according to section 2.2.4.

#### 2.3.5 Fine root emergence and turnover

Absorptive fine roots were represented as short-lived, non-lignifying lateral roots emerging exclusively from living structural roots. Their initiation follows the same successor-based branching framework as structural roots, with age-dependent branching probabilities assigned to each structural subtype (section 2.3.2; Supplementary Methods S1).

Fine roots were implemented as terminal modules: they do not branch further, undergo secondary growth, or lignify, and do not contribute to the long-term structural framework. Axial elongation follows a negative exponential function until a subtype-specific maximum length (typically 5 cm) is reached, consistent with empirical observations (Mapfumo et al. 1994).

Turnover of fine roots is linked to developmental status and parental survival rather than explicit age-based lifetimes. Fine roots are removed at the onset of winter or when their parent structural root dies.

#### 2.3.6 Developmental status transitions (suberisation and lignification)

To represent anatomical and functional transitions along grapevine roots, we implemented three categories of development status: unsuberised, suberised, and lignified tissue states. This formulation follows anatomical descriptions of grapevine roots reporting strong declines in radial hydraulic conductivity (*L*_pr_, *m s^−^*^1^ *MPa^−^*^1^) with increasing distance from the apex due to suberisation and secondary growth (Gambetta et al. 2013).

In the model, each root segment is assigned a developmental status based on segment age. These transitions regulate winter survival, *L*_pr_, and eligibility for secondary growth, thereby linking anatomical maturation to hydraulic function and long-term structural dynamics. For instance, unsuberised are removed at the onset of winter. Threshold definitions and implementation details are provided in Supplementary Methods S5.

#### 2.3.7 Hydraulic parameterisation

Root hydraulic properties were parameterised by coupling root xylem anatomy and developmental status to empirically constrained axial conductances (*K*_h_, *m*^4^ *s^−^*^1^ *MPa^−^*^1^) and radial conductivities (*L*_pr_, *m s^−^*^1^ *MPa^−^*^1^). *K*_h_ of structural roots was calculated as a function of root diameter using genotype-specific diameter–conductance relationships derived for grapevine rootstocks (Fichtl et al. 2024). These relationships were applied dynamically during simulation.

*L*_pr_ was assigned according to developmental status (unsuberised, suberised, lignified), using representative values from grapevine measurements (Cuneo et al. 2018; Gambetta et al. 2013). Consequently, *K*_h_ increases with diameter and age, whereas *L*_pr_ declines with tissue maturation.

*K*_h_ and *L*_pr_ were integrated into the CPlantBox hydraulic solver (Vanderborght et al. 2024a,b) to compute segment-level uptake and whole-root-system conductance (*K*_rs_, *m*^4^ *s^−^*^1^ *MPa^−^*^1^). Parameter values and implementation details are provided in Supplementary Methods S6 and Table S2.

#### 2.3.8 Estimation of maximum attainable structural root length

Maximum attainable structural root lengths (*L*_max_, *cm*) could not be measured directly from excavations. We therefore estimated order- and rootstock-specific maximum lengths from the temporal development observed in the young-vine dataset.

For each rootstock and structural order, the longest observed roots at successive excavation dates were related to root age since order-specific emergence. These trajectories were used to infer asymptotic maximum lengths for each order. The resulting *L*_max_ values were applied as order-specific constraints in the age-dependent growth module. Details of the fitting procedure are provided in Supplementary Methods S7.

#### 2.3.9 BEDD-dependent growth potential

To couple empirically derived age-dependent growth decline with climatic forcing, root elongation was converted from an age-driven process (see Eqn. 2) to a process driven by biologically effective degree days (BEDD, see Supplementary Methods S2). The parameter *β* was calibrated for each root type such that BEDD-based elongation trajectories reproduced the empirically derived age-dependent decline in growth potential. In this formulation, age-dependent damping emerges implicitly from thermal-time saturation rather than being imposed explicitly. Parameter estimation details are provided in Supplementary Methods S8.

#### 2.3.10 Time units and numerical time stepping

Plant growth simulations were performed in BEDD: BEDD accumulate daily heat units above a base temperature *T*_base_ = 10*^◦^*C and below an upper effectiveness threshold *T*_cap_ = 19*^◦^*C. For a given day *d* with mean air temperature *T_d_*, daily BEDD were computed as

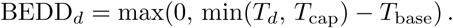

In this study, one biological year corresponded to a fixed internal time step of

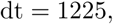

representing the long-term mean annual BEDD at the study site. All age-related quantities, including root age, structural root lifetimes, and branching thresholds, were expressed in this common unit. Survival and dormancy transitions were evaluated once per annual time step.

Several processes were simulated stochastically. Structural root lifetimes were drawn from order-specific Weibull distributions at organ creation. Lateral identity (structural versus fine root) was assigned probabilistically according to age-dependent branching probabilities. Segment orientation followed a probabilistic mixture of tropistic responses using genotype-specific weights, generating replicate-level variability in three-dimensional architecture.

#### 2.3.11 Simulation initialisation and output extraction

Perennial simulations were initialised from empirically measured one-year-old grapevine root systems. For each rootstock genotype (‘101-14’, ‘SO4’, ‘R110’), genotype-specific RSML files (see section 2.4) from the vineyard establishment experiment (Fichtl et al. 2024, 2025) were imported, and stem and initial (static) roots (orders 0–1) were mapped to structural subtypes in CPlantBox. This ensures that all long-term simulations started from realistic, field-derived architectures of planted rooted cuttings rather than synthetic seedlings.

Root development was simulated over *N* = 50 years with eight stochastic replicates per genotype using the annual BEDD-based time step. Deterministic parameter sets were identical within genotypes; variability arose from stochastic branching, tropism, and mortality processes.

Model outputs were summarised annually. For each root order, we extracted root numbers, length distributions, and age distributions. Simulated age structures were compared with empirical age distributions represented as Gaussian kernel density estimates (KDEs), using mean and standard deviation across vines as reference envelopes.

#### 2.3.12 Standard uptake fractions and *α*–*ω* upscaling

The hydraulic structure of the root system was characterised using the standard uptake fraction (SUF, dimensionless) (Couvreur et al. 2012), computed after each annual growth step from the coupled root hydraulic model. SUF represents the fraction of total uptake contributed by a root segment (or soil layer) under uniform soil water potential and satisfies

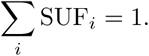

Segment-wise radial inflow was normalised by total transpiration under equilibrium soil conditions using the Meunier hydraulic solver (Couvreur et al. 2012; Meunier et al. 2017). SUF values were aggregated to depth layers to obtain depth-discrete profiles.

Depth-resolved SUF profiles were interpreted using the mechanistic *α*–*ω* framework (Vander-borght et al. 2024a,b). The layer-specific supply ratio *α* was defined as

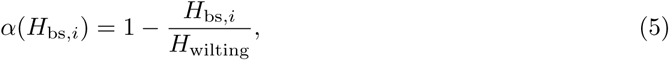

with *H*_bs_ (*cm*) the water head of the bulk soil at layer *i* and *H*_wilting_ (*cm*) the plant’s wilting point. The whole-root-system supply ratio as the SUF-weighted average

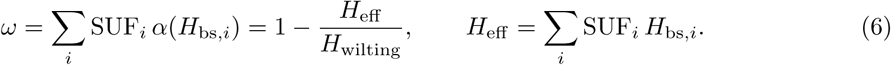

These quantities provide a mechanistic upscaling from explicit hydraulic architecture to profile-scale water supply metrics.

#### 2.3.13 Hydraulic scenarios

Two idealised precipitation regimes were imposed (Table 1). In the “continental” scenario, initial soil total potential was set to *−*5000 *cm*, assuming 50% winter recharge of plant-available water (Tron et al. 2015), followed by moderate rainfall during the growing season (50 mm per month, April–October). In the “Mediterranean” scenario, initial total potential was *−*330 *cm* (full winter recharge) with no precipitation during the growing season.

**Table 1:**
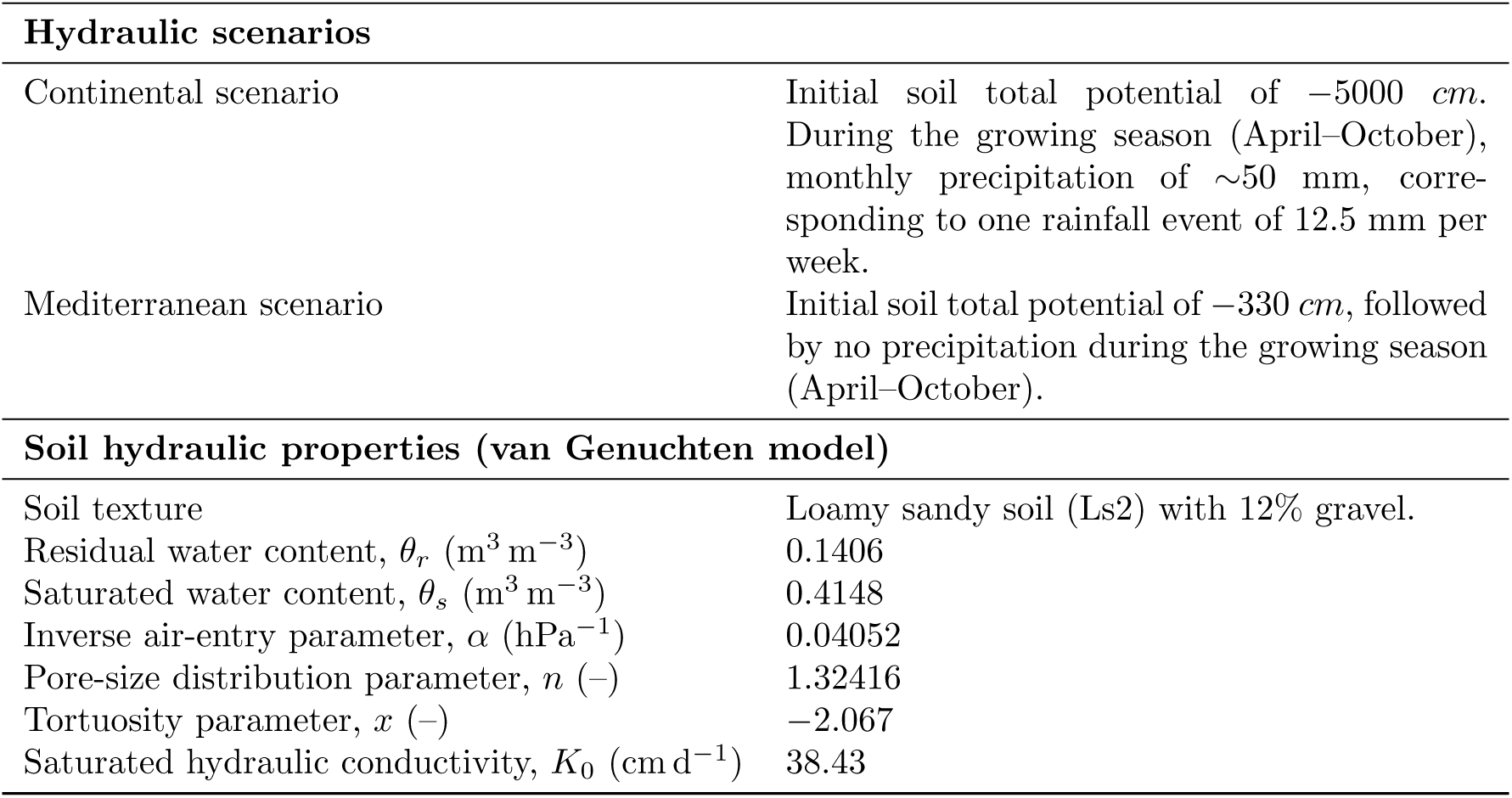
Hydraulic scenarios and soil hydraulic parameters used in the perennial model simulations. Hydraulic scenarios represent idealised precipitation regimes, and soil hydraulic properties are based on measurements from the vineyard site where 48-year-old vines were excavated.

| Hydraulic scenarios |  |
| --- | --- |
| Continental scenario | Initial soil total potential of $-5000$ cm. During the growing season (April–October), monthly precipitation of $\sim 50$ mm, corresponding to one rainfall event of 12.5 mm per week. |
| Mediterranean scenario | Initial soil total potential of $-330$ cm, followed by no precipitation during the growing season (April–October). |
| Soil hydraulic properties (van Genuchten model) |  |
| Soil texture | Loamy sandy soil (Ls2) with 12% gravel. |
| Residual water content, $\theta_r$ ($\text{m}^3 \text{m}^{-3}$) | 0.1406 |
| Saturated water content, $\theta_s$ ($\text{m}^3 \text{m}^{-3}$) | 0.4148 |
| Inverse air-entry parameter, $\alpha$ ($\text{hPa}^{-1}$) | 0.04052 |
| Pore-size distribution parameter, $n$ (–) | 1.32416 |
| Tortuosity parameter, $x$ (–) | $-2.067$ |
| Saturated hydraulic conductivity, $K_0$ ($\text{cm d}^{-1}$) | 38.43 |

Soil hydraulic properties were described using the van Genuchten–Mualem model parameterised for the vineyard soil (loamy sandy soil, Ls2, 12% gravel) (Linsenmeier et al. 2008), with parameters taken from published tables (Wessolek et al. 2009) and held constant across simulations.

Potential transpiration was prescribed as a truncated sinusoidal daily function scaled to 2 L h*^−^*^1^. The precipitation was likewise converted to hourly inputs. A no-flux upper boundary (except rainfall) and free-drainage lower boundary were imposed. Hydraulic simulations used a temporal resolution of 1 h and a vertical discretisation of 1 cm.

Plant water potential was computed at the end of each simulation using a Neumann boundary condition at the root collar based on the final transpiration rate.

#### 2.3.14 Data-guided parameterisation of demographic and branching processes

Model parameterisation followed a data-guided approach rather than formal optimisation. The objective was to identify parameter sets reproducing key empirical features of grapevine root system architecture and demography across establishment and mature stages.

Order-specific survival parameters were derived directly from empirical age distributions. Weibull shape and scale parameters were estimated from Kaplan–Meier survival curves of ring-counted basal ages of “main”, “sub”, and “subsub” roots and transferred unchanged into the model to define stochastic lifetimes.

Branching parameters (age-dependent probabilities and maximum lateral numbers) were constrained using young-vine architectural observations and adjusted to remain consistent with mature demographic patterns.

#### 2.3.15 Derivation of rootstock-specific parameter sets

Rootstock-specific parameter sets combined (i) core architectural and hydraulic parameters inherited from CPlantBox, (ii) perennial extensions governing age-dependent demography and growth, and (iii) genotype-specific traits derived from young-vine observations (Supplementary Table S4).

Core parameters describe elongation, branching, tropisms, and axial and radial hydraulic properties. Perennial extensions include age-dependent survival, secondary radial growth, dormancy-related root loss, and thermal-time-dependent damping of elongation. Perennial parameters were derived from the empirical analyses presented above, including age distributions, age–diameter relationships, and reconstructed lifetime thermal exposure.

Rootstock-specific parameterisation was restricted to traits constrained by the young-vine dataset, including maximal root length (*L*_max_), branching behaviour, and axial hydraulic conductance (*K*_h_) functions. Age-dependent survival, secondary growth rates, and dormancy-related turnover rules were applied uniformly across rootstocks.

Parameter sets were evaluated using a multi-criteria approach. Simulations were retained if they reproduced (i) root numbers and total root length during the first two years after planting, (ii) fine-to-structural root length ratios in mature systems, and (iii) structural root age distributions observed in 48-year-old vines.

#### 2.3.16 Model evaluation

Model performance was assessed by comparing simulated and observed root system properties using three descriptors: (i) survival probability distributions of structural roots, (ii) order-specific root numbers over system age, and (iii) the fine-to-total root length ratio.

Simulated age distributions were compared with empirical kernel density estimates (KDEs) derived from mature vine excavations. Order-specific root numbers were evaluated against observed counts from young and mature 3D datasets.

Because fine-root length was not quantified in mature excavations (see section 2.1.2), the simulated fine-to-total root length ratios were compared with literature-based estimates for mature grapevines. Lanari et al. (2025) reported fine-to-total root length ratios ranging from 81-87% (diameter threshold: 1 *mm*), while Steng et al. (2026) observed a mean ratio around 91% (diameter threshold: 3 *mm*), although the latter study reported a standard deviation sufficiently large for the ratio to reach values as low as 65%. We therefore set a rough estimate of 80% for our objective ratio.

### 2.4 Data Handling, Analysis and Visualisation

Post-processing and analyses of empirical and simulated data were conducted using R (v4.5.1; R Core Team (2024)), using the graphical user interface RStudio (v2025.09; RStudio Team (2020)). Empirical root architecture data from young and mature grapevines were processed from RSML (RootSystemML) files (Lobet et al. 2015). RSML is a file format based on the XML standard, created to store root architectural data. Root orders, lengths, and branching information were derived directly from the digitised geometries. Survival analyses of structural roots were performed using the survival (Therneau 2026) and survminer (Kassambara et al. 2025) packages. Age-dependent growth damping functions and secondary growth relationships were fitted using nonlinear least squares with the minpack.lm package (Elzhov et al. 2023). Simulation outputs from CPlantBox were written to columnar Parquet format for quantitative analysis in R, while VTP (VTK PolyData) files were generated for three-dimensional visualisation. All two-dimensional figures were generated using ggplot2 and related packages from the tidyverse ecosystem (Wickham 2016). Three-dimensional visualisations of simulated root systems and hydraulic states were produced using ParaView (v5.12.0; Ahrens et al. (2005)).

## 3 Results

### 3.1 Empirical Root System Data from Young and Mature Grapevines

We combined three-dimensional root system data from young grapevines (1–2 years after planting) with excavations of mature, 48-year-old vines.

In mature vines, lignified root architecture was restricted to a limited set of branching orders beyond the initial static roots, comprising “main” roots, first-order laterals (“sub”), and higher-order laterals (“subsub”) (Supplementary Figure S1). No lignified roots of higher order or unsuberised fine roots were observed in the mature excavations. The absence of higher order structural roots could be explained by the partial senescence of the root system. On the other hand, the absence of fine roots was assumed to be caused by the fact that the root system were excavated in winter (see section 2.1.2).

These observations were completed with literature data and used to define the functional root typology and to parameterise order-specific survival, branching, and fine-root turnover in the perennial model.

### 3.2 Root Age Demography and Age-Dependent Growth Regulation

Annual growth rings were assessed as age indicators in grapevine roots. Transverse sections of young roots of known age (1–4 years after planting) showed distinct, countable concentric rings (Figure 3A). Across all three rootstock genotypes, root diameter increased approximately linearly with age during early development (Figure 3B).

**Figure 3:**
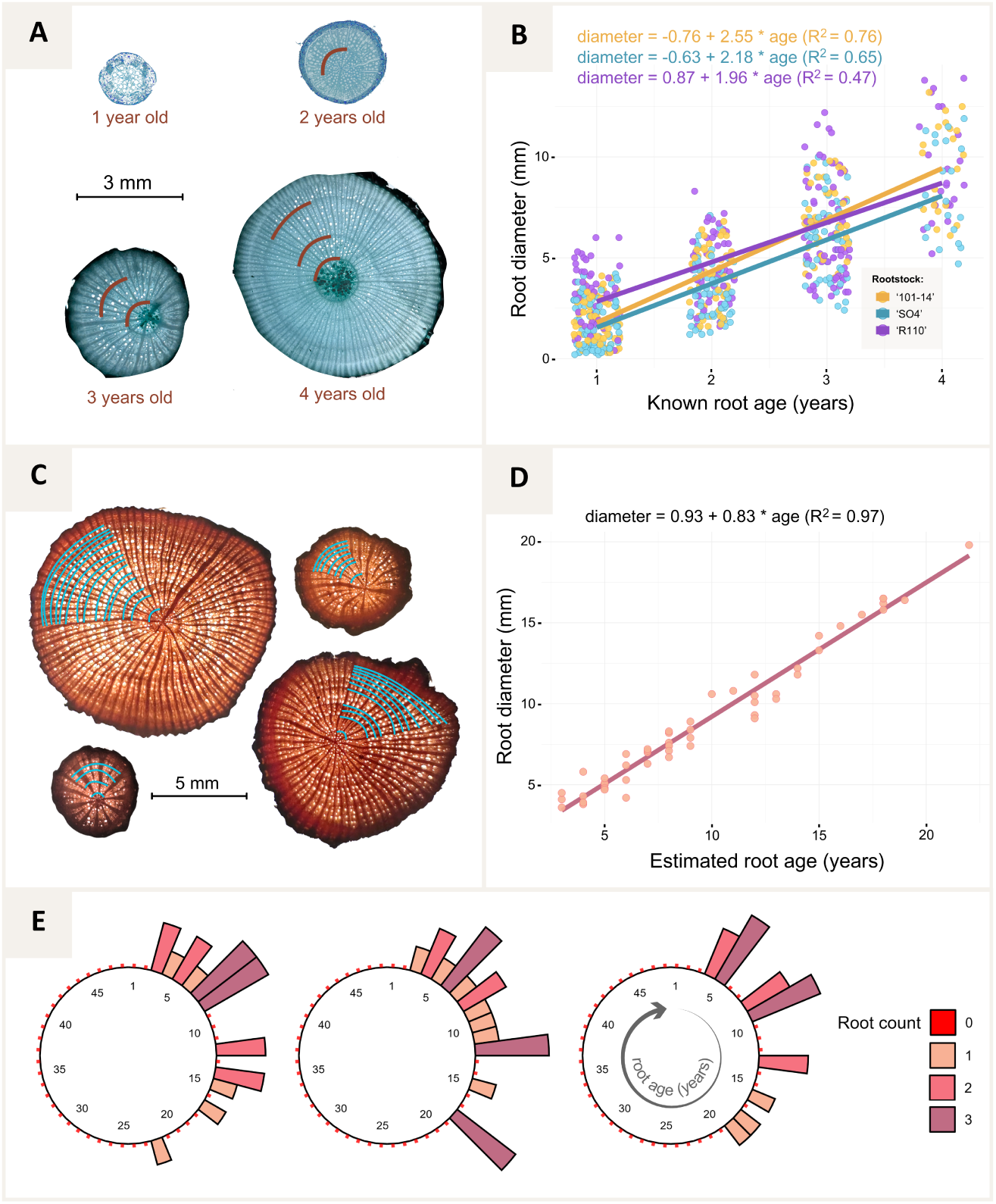
Validation of root age estimates and woody root age distributions in mature grapevines. (A) Transverse sections of young roots of known age (1–4 years) showing distinct, countable annual growth rings (red arcs). (B) Linear relationships between root diameter and known age for young roots of three rootstock genotypes, demonstrating approximately linear radial growth during early years. (C) Representative cross-sections of trunk-borne roots from 48-year-old vines with ring counts indicated (blue arcs). (D) Linear regression of basal diameter on ring-counted age for all “main” roots from mature vines (*n* = 54); diameter increases nearly linearly with age (equation and *R*^2^ shown). (E) Circular histograms of basal age distributions of trunk-borne roots for each excavated mature vine. Bars represent the number of roots in each 1-year age class (colour intensity indicates root count), revealing coexisting cohorts of young and old structural roots and a right-skewed age distribution extending beyond 20 years.

The same approach was applied to mature vines. Trunk-borne woody roots from 48-year-old plants exhibited well-defined ring structures over extended time spans (Figure 3C). Ring-counted basal ages ranged from 3 to 23 years (mean 9.5 years). Basal diameter increased linearly with age (R² = 0.97; Figure 3D).

Age distributions of trunk-borne roots showed multi-cohort structures in all excavated vines (Figure 3E). Individual plants contained young, intermediate-aged, and old woody roots, with right-skewed age distributions extending beyond 20 years. Most roots belonged to intermediate age classes (approximately 2–10 years), whereas very young and very old roots were less frequent.

#### 3.2.1 Secondary radial growth dynamics

Basal root diameter increased nearly linearly with ring-counted age in mature vines (Figure 3D), following Eq. 4:

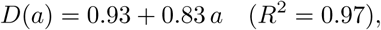

where *D* is basal diameter (mm) and *a* root age (years). The slope corresponds to an average secondary thickening rate of 0.83 mm yr*^−^*^1^ in diameter (0.415 mm yr*^−^*^1^ in radius under symmetric growth).

#### 3.2.2 Order-specific survival probabilities and structural root turnover

Order-specific survival functions were reconstructed for the three structural root classes (“main”, “sub”, “subsub”). For “main” roots, age was obtained directly from basal ring counts, whereas ages of first- and higher-order laterals were inferred from the age–diameter relationship (Figure 3D).

Kaplan–Meier estimators showed a decline in survival probability with increasing age in all three orders, with clear order-specific contrasts (Figure 4). “Main” roots exhibited the slowest decline, with non-zero survival probabilities extending beyond 20 years. Sub roots declined more steeply and were rare beyond approximately 14 years, while subsub roots were concentrated within the first decade and rarely exceeded 8 years.

**Figure 4:**
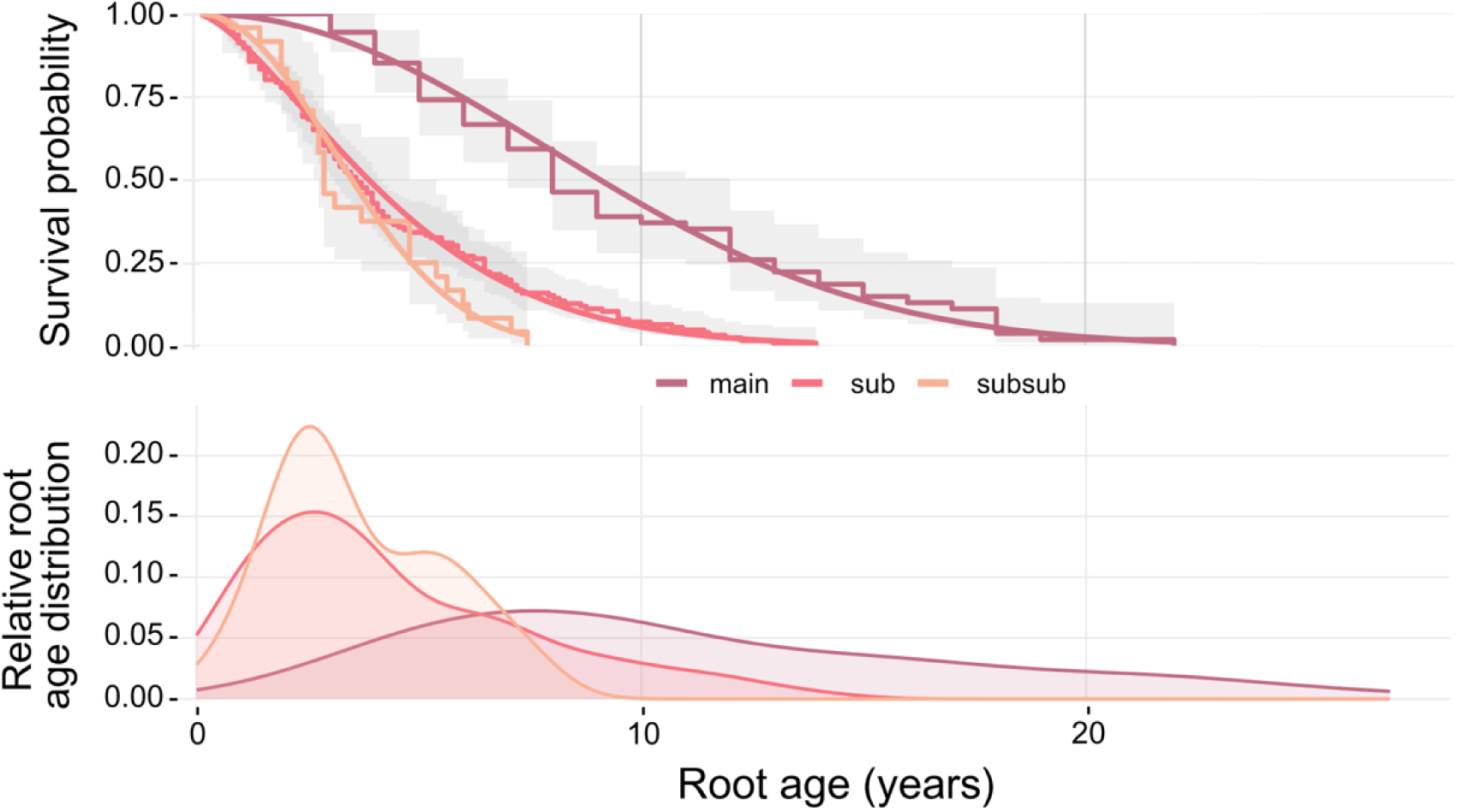
Order-specific survival and root age distribution of woody grapevine roots. (Top) Kaplan–Meier estimates (step curves with grey confidence bands) and fitted Weibull survival functions (smooth coloured lines) for “main”, “sub” and “subsub” roots based on basal ages of living structural roots in 48-year-old vines. For each order, the Weibull fit was truncated at the maximum age observed in the data (“main”: 23 years; “sub”: 14 years; “subsub”: 8 years), and survival probabilities were set to zero beyond this age. (Bottom) Empirical age distributions of the corresponding root orders derived from the dendrochronological dataset from mature vines. which were used to parameterise stochastic structural root mortality in the perennial model (Supplementary Materials S3; Table S1).

Parametric Weibull models closely matched the observed Kaplan–Meier curves across all orders (Figure 4). Fits were truncated at the maximum ages observed in the empirical data (23 years for “main”, 14 years for “sub”, and 8 years for “subsub” roots).

The resulting shape (*k*) and scale (*λ*) parameters define age-dependent annual survival probabilities,

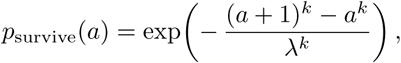

#### 3.2.3 Maximum root length and thermal-time–dependent damping of elongation potential

Order-specific maximal root lengths (*L*_max_) were estimated from young grapevines, as mature excavations captured only truncated terminal lengths. Across genotypes, “main” roots reached asymptotic lengths of 630–790 cm, whereas lateral structural roots were substantially shorter (“sub”: 90–120 cm; “subsub”: 50–60 cm; Supplementary Table S3).

Relative realised root length increased with accumulated biologically effective degree days (BEDD) following an exponential saturation,

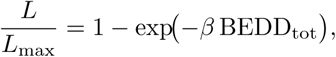

where *β* describes thermal sensitivity of elongation. Thermal saturation occurred most rapidly in “main” roots, whereas elongation of sub and subsub roots approached their maxima more gradually (Figure 5).

**Figure 5:**
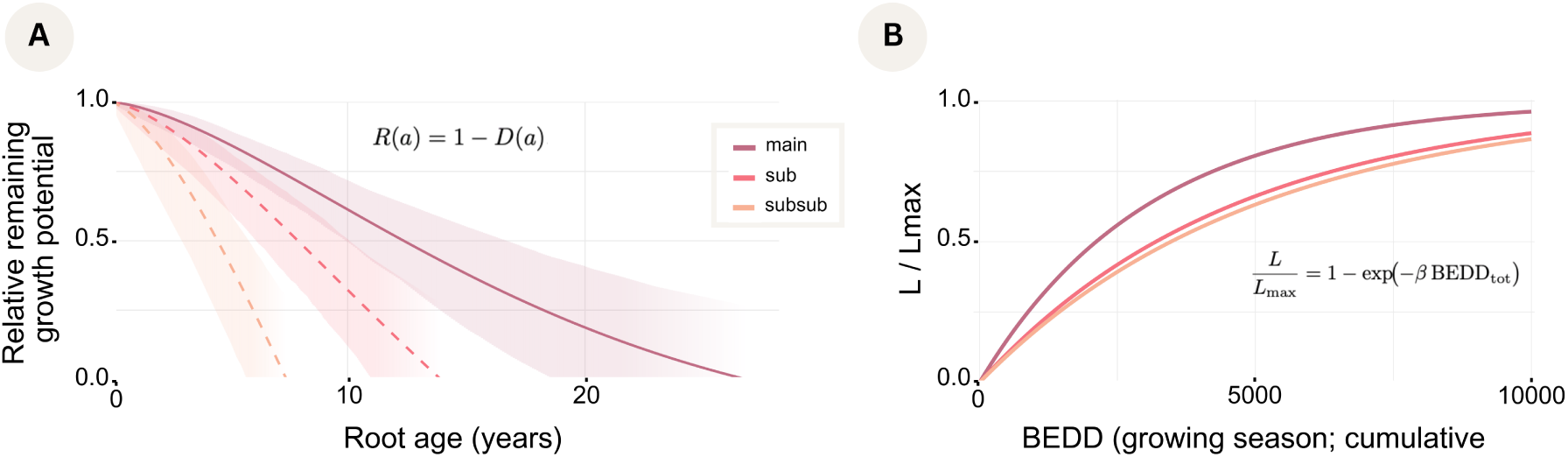
Age-based remaining growth potential and its thermal-time implementation in the perennial root model. (A) Empirically derived remaining growth potential of structural roots as a function of root age. Curves are shown for “main”, “sub” and “subsub” roots; shaded areas indicate uncertainty derived from parameter resampling. Functions are truncated at the maximum basal ages observed for each root order. (B) Continuous thermal-time growth formulation implemented in the model, expressed as the realised length fraction. Order-specific *β* parameters reproduce the empirically inferred age-dependent decline in remaining growth potential, while *L*_max_ constrains absolute elongation.

### 3.3 Model Validation

#### 3.3.1 Model validation across structural and functional scales

Simulated root system dynamics were evaluated against empirical observations at multiple structural levels.

Simulated survival probability distributions reproduced the observed age-dependent survival patterns across all three structural orders (Supplementary Figure S2). Agreement was highest for “main” and first-order lateral roots, whereas deviations increased for higher-order woody roots.

Simulated root number trajectories captured order-specific patterns of root abundance during development (Supplementary Figure S3). Agreement was strongest for low-order structural roots, while discrepancies increased with root order. Across both validation metrics, model performance varied primarily with root order rather than genotype.

#### 3.3.2 Validation of the fine-to-total root length ratio

The ratio of fine-to-total root length ratios was evaluated for the young grapevine by defining all roots as fine roots which have a length below 5 cm and a diameter below 0.93 mm (which is the minimum diameter of a structural root, see section 3.2.1). Because the mature 3D excavations quantified structural roots only (see section 2.1.2), fine-root length could not be validated directly for older grapevines. Based on the literature, a fine-to-total root length ratio of approximately 80% was assumed for mature grapevines (see section 2.3.16).

Across all rootstocks, simulated fine-to-total root length ratios increased rapidly during early development and stabilised after approximately 10–12 years (Figure 6). In the mature phase, simulations converged towards stable values close to the literature benchmark, with modest but consistent differences among rootstocks. Inter-run variability was low after establishment.

**Figure 6:**
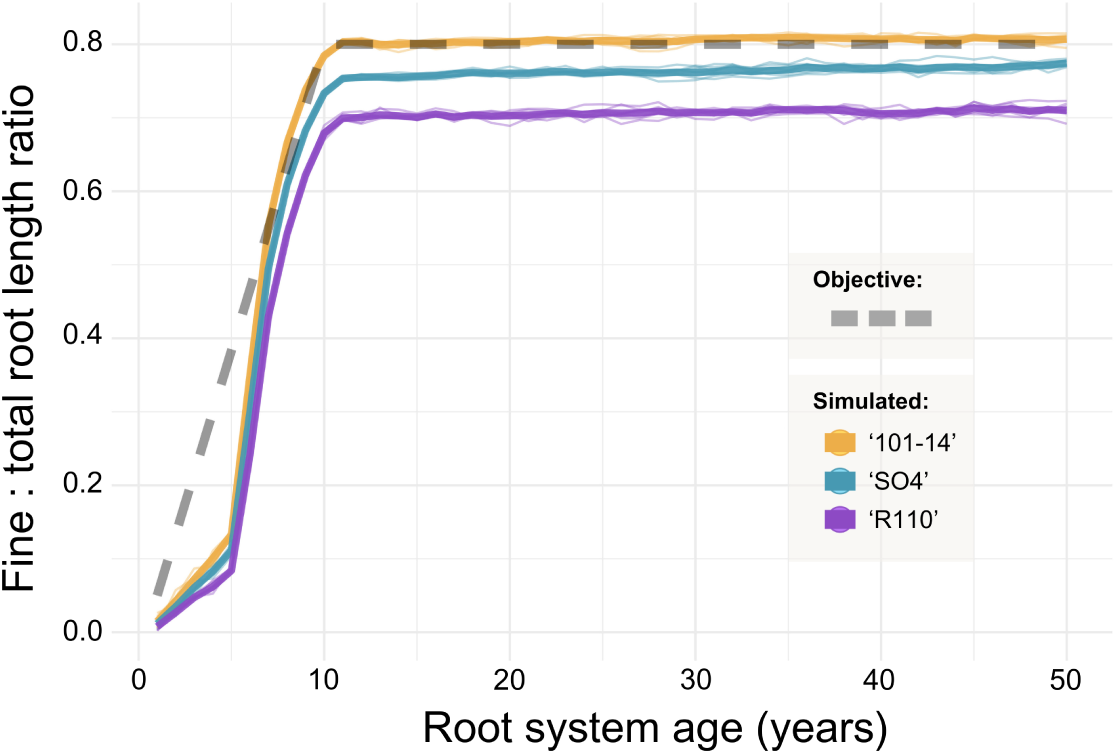
Validation of simulated fine-to-total root length ratios. Simulated trajectories are shown for the three grapevine rootstock calibrations, obtained from the experimental data. Thin coloured lines represent individual simulation runs, and thick lines indicate genotype-specific means. The dashed grey line denotes the objective values that are based on experimental observations and a literature review (see section 3.3.2).

### 3.4 Emergent Architecture and Functional Consequences

#### 3.4.1 Perennial 3D root system simulations

Three-dimensional root system development was simulated over multiple decades. Figure 7 shows representative simulations for three rootstocks (‘101-14’, ‘SO4’, and ‘R110’) at 2, 5, 10, 25, and 50 years after planting.

**Figure 7:**
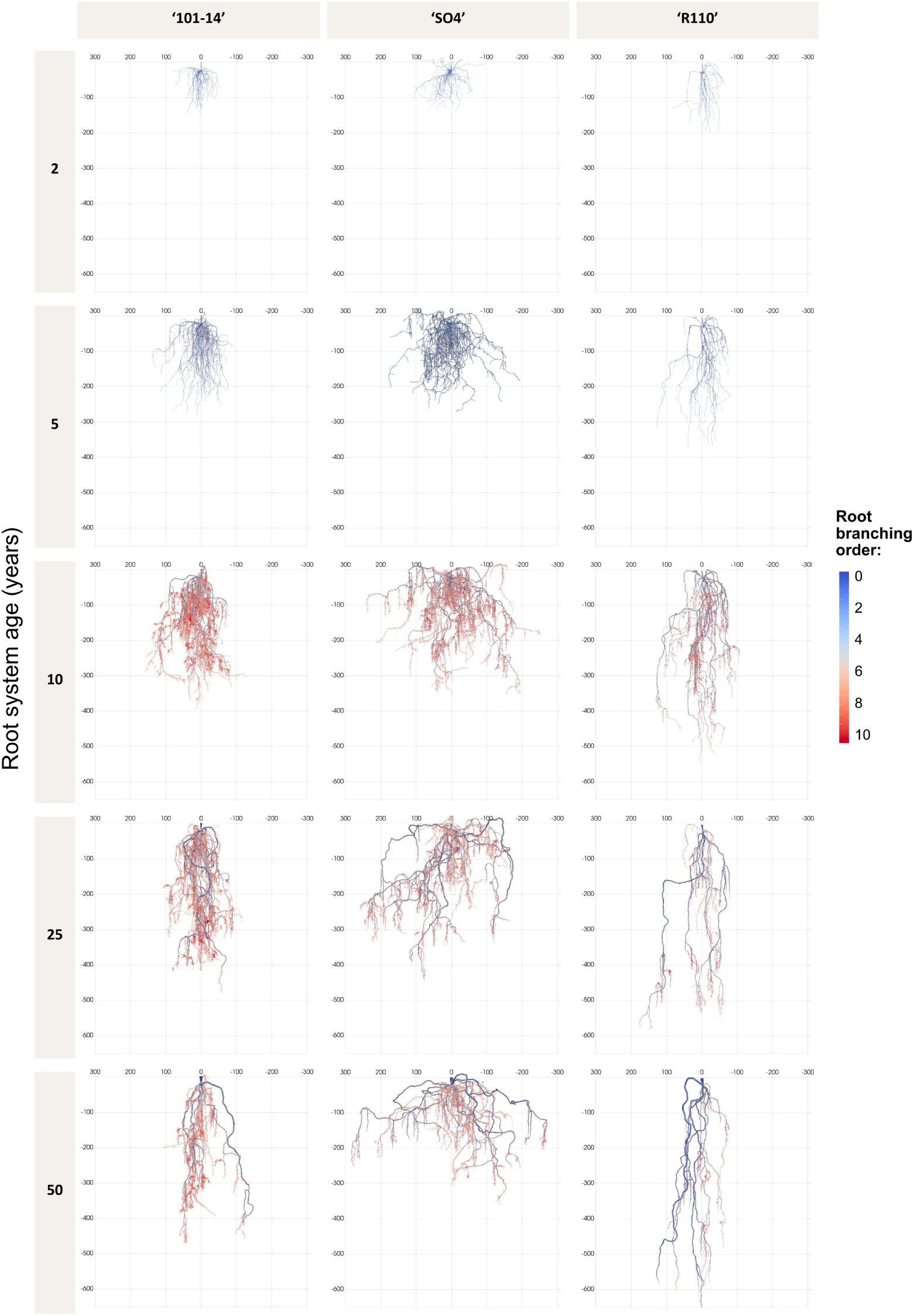
Examples of simulated grapevine root systems for three rootstocks (columns: ‘101-14’, ‘SO4’, and ‘R110’) shown at increasing root system ages (rows: 2, 5, 10, 25, and 50 years). Individual roots are coloured by branching order, ranging from low-order structural axes (blue) to higher-order laterals and fine roots (red).

Across all rootstocks, early development was characterised by rapid establishment of a small number of dominant structural axes, followed by extensive proliferation of lateral and fine roots during the first decade. With increasing age, root systems showed reduced net expansion of major axes, while higher-order laterals and fine roots continued to turn over.

Distinct rootstock-specific architectures were maintained over time. ‘SO4’ developed a laterally extensive root system, ‘R110’ showed a more vertically oriented architecture dominated by deep “main” roots, and ‘101-14’ displayed intermediate lateral spread and depth. After approximately two decades, overall architectural patterns changed only gradually, with subsequent modifications occurring primarily at local scales.

#### 3.4.2 Age-dependent standard uptake fractions

Vertical profiles of standard uptake fraction (SUF) were analysed for three rootstocks across increasing root system ages (Figure 8). Mean SUF values per 10-cm soil layer were derived from replicated simulations (n = 8 per genotype).

**Figure 8:**
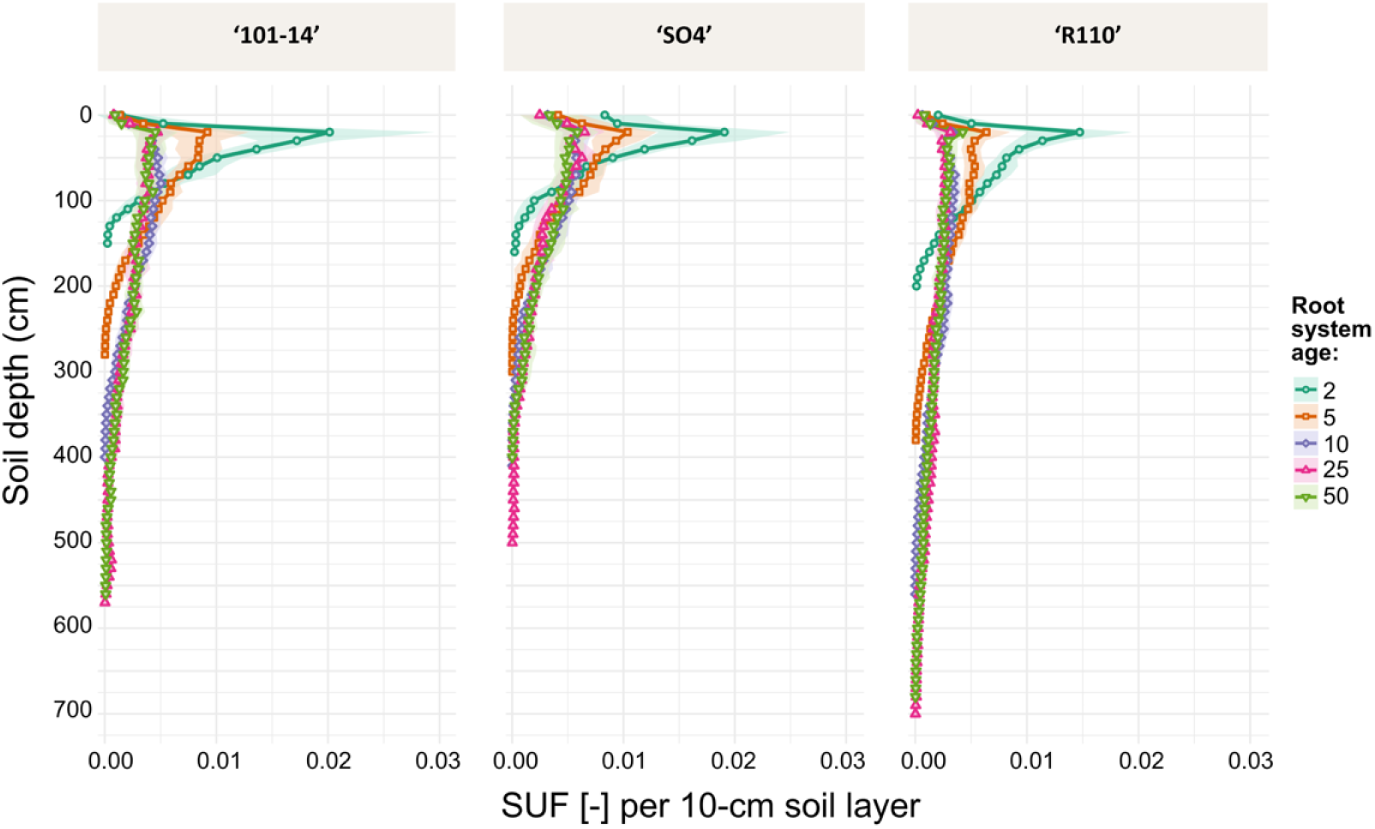
Age-dependent vertical profiles of standard uptake fraction (SUF) in simulated perennial grapevine root systems. Mean profiles of fractions per 10-cm soil layer are shown as a function of soil depth for three rootstocks (‘101-14’, ‘SO4’, and ‘R110’; columns) at increasing root system ages (2, 5, 10, 25, and 50 years). Coloured lines represent the mean SUF across replicate simulations, with shaded ribbons indicating ± SD among simulation runs (n = 8 for each genotype). For each time point and root system, the SUF sums up to one. Higher values of SUF indicate the areas where the root system would take up the most water under a soil water potential at equilibrium.

In all rootstocks, young root systems (2–5 years) showed strongly surface-weighted SUF profiles, with most uptake potential concentrated in shallow soil layers. With increasing age, SUF profiles extended progressively to greater depths, accompanied by a relative decline in uptake contribution from near-surface layers. Most vertical redistribution occurred during the first decade after planting, after which SUF profiles changed only gradually.

Distinct rootstock-specific uptake patterns were maintained in mature systems. ‘SO4’ displayed comparatively broad SUF distributions across intermediate soil depths, whereas ‘R110’ exhibited more depth-concentrated profiles. ‘101-14’ showed an intermediate pattern with moderate surface uptake and progressive deepening over time.

#### 3.4.3 Architecture-driven hydraulic responses under contrasting climates

Water uptake and transpiration were simulated across development under two simplified climate scenarios (“continental” and “mediterranean”). Representative 50-year-old root systems for three rootstocks (‘101-14’, ‘SO4’, and ‘R110’) are shown in Figure 9, with roots coloured by simulated xylem pressure head.

**Figure 9:**
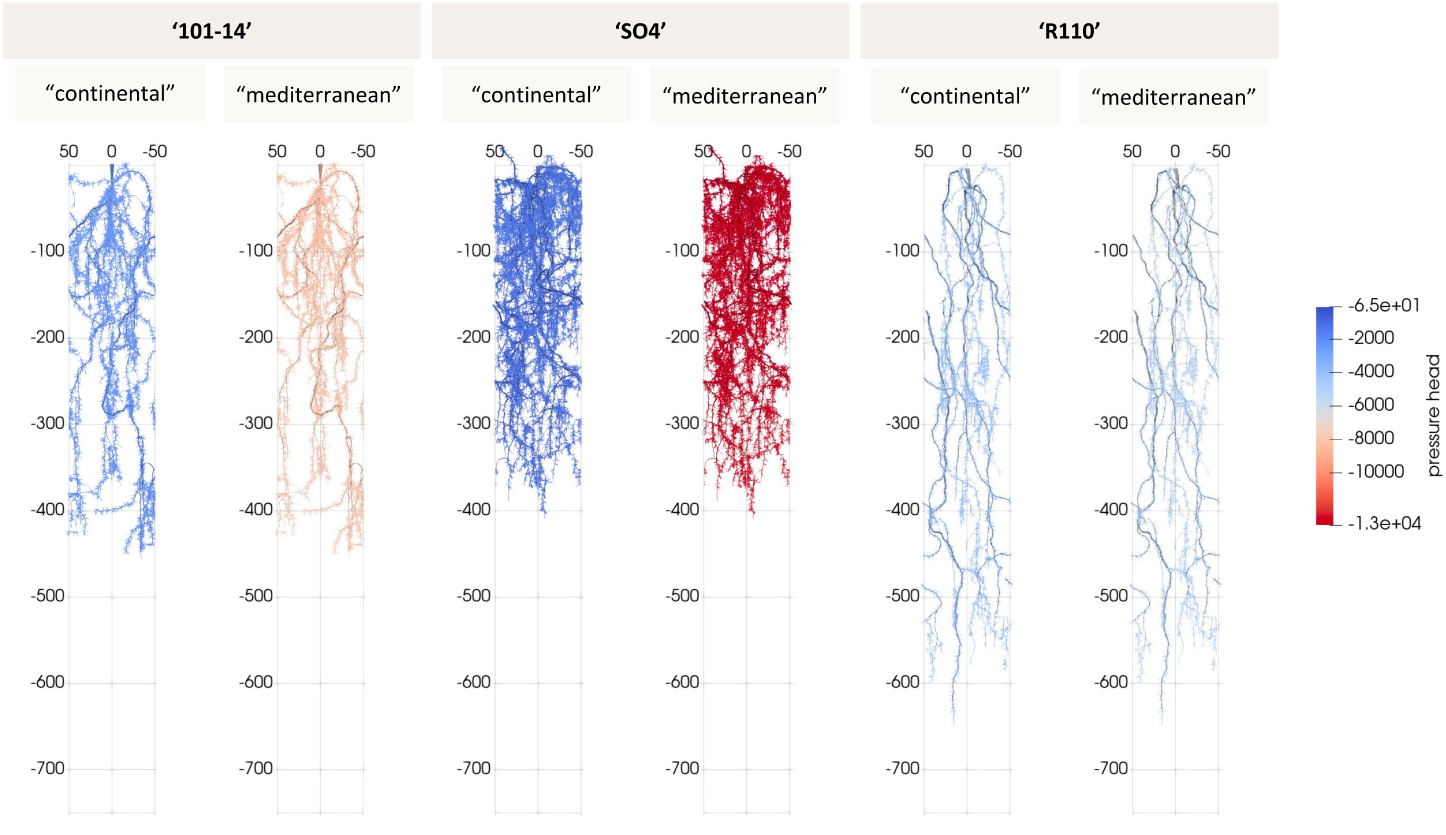
Representative simulated root systems (50 years old) for three grapevine rootstocks (‘101-14’, ‘SO4’, and ‘R110’) under two contrasting climatic scenarios (columns within each rootstock: “continental” and “mediterranean”). Roots are coloured by simulated xylem pressure head (cm), with blue indicating higher (less negative) and red indicating lower (more negative) water potentials. All simulations depict the effective below-ground soil volume available to a single vine under a typical cropping system with a planting distance of 1 m between vines.

Across all grapevines, cumulative transpiration increased with root system age. In young systems, genotype-specific differences were apparent under mediterranean conditions. ‘SO4’ and ‘101-14’ showed reductions in cumulative transpiration relative to imposed potential demand during the growing season, particularly from mid-season onwards. In contrast, ‘R110’ achieved the prescribed daily transpiration rate across simulated ages and under both climate scenarios.

With increasing age, these differences diminished. By 50 years, all genotypes reached the target transpiration demand under both scenarios. Differences in simulated xylem pressure head remained evident. ‘R110’ exhibited comparatively stable xylem water potentials across scenarios. ‘SO4’ and ‘101-14’ showed stronger scenario-dependent variation, with marked declines in xylem pressure head under mediterranean conditions. Among these, ‘SO4’ operated at lower water potentials than ‘101-14’.

## 4 Discussion

### 4.1 Modelling of the vine root system structure

This study introduces a perennial root architecture modelling framework that extends CPlantBox from annual to multi-decadal simulations by explicitly representing age-structured demography, dormancy and secondary growth. A central advance is the integration of empirically derived woody-root age distributions, survival functions and age-dependent growth damping into the model core, so that stochastic root emergence, elongation and mortality are constrained by observations rather than prescribed heuristics. By distinguishing long-lived structural and short-lived absorptive roots as functional types independent of branching order, the framework decouples architectural hierarchy from function and captures gradients in longevity, anatomy and hydraulic role. A BEDD-driven dormancy module links root development to climatic forcing, and coupling developmental status with axial conductances (*K*_h_) and radial conductivities (*L*_pr_) provides a hydraulically explicit representation of perennial RSA. Together, these components enable lifelong simulations in which architecture, demography and function arise from empirically grounded processes rather than static lifespan assumptions.

### 4.2 Biological findings

Beyond the modelling advance, our empirical analyses provide a clear biological message: woody roots in grapevine do not form a permanent architectural scaffold, but undergo continuous turnover over decadal timescales. Dendrochronological analyses revealed pronounced multi-cohort age structures in all excavated mature vines, with coexistence of young and old woody roots and right-skewed age distributions consistent with continuous recruitment and age-dependent mortality. Although the root system could not be monitored continuously over the 48-year experiment, earlier studies covering the 1990–2006 and 2017–2018 periods (Friedel et al. 2020; Linsenmeier 2007) did not find any indication of root system collapse due to environmental stresses. Moreover, the shape parameter of the fitted Weibull survival function was similar across root orders even though the scale parameter varied: structural (main) roots therefore appear to senesce according to the same underlying pattern of ageing as higher-order roots, only on a slower timescale, rather than dying off abruptly from an acute stress event. These findings challenge the implicit assumption that lignified structural roots persist for the lifetime of the plant (Freschet et al. 2021). Instead, perennial root systems function as demographically dynamic populations in which woody transport pathways are continuously replaced. This provides rare empirical constraints on decadal woody-root turnover, a process long recognised as important for perennial persistence but difficult to quantify because root longevity and mortality remain methodologically challenging to assess (Comas et al. 2000).

### 4.3 Modelling of the vine root system water uptake

Embedding age-dependent survival, elongation damping and secondary growth into a unified framework clarifies how demographic processes translate into long-term architecture and hydraulic function. In the simulations, age-structured turnover combined with progressive damping of elongation stabilised mature architectures while maintaining continuous renewal of higher-order woody roots. These architectural dynamics were reflected in vertical patterns of standard uptake fraction (SUF), shifting from shallow, establishment-phase uptake to deeper and more evenly distributed uptake profiles in mature systems. Hydraulic scenario simulations further illustrate how perennial root development modulates whole-plant water supply capacity over decades and interacts with climatic forcing in a genotype-specific manner. Hydraulic capacity increased with root system age in all genotypes, such that mature systems sustained the prescribed potential transpiration under both climatic scenarios. However, the hydraulic state underlying this compensation differed among genotypes. Under continental conditions, ‘SO4’ and ‘101-14’ operated at higher (less negative) xylem water potentials than ‘R110’, whereas under mediterranean conditions both genotypes exhibited markedly reduced xylem water potentials, with ‘SO4’ consistently more negative than ‘101-14’.

In contrast, ‘R110’ maintained comparatively stable xylem water potentials across scenarios while sustaining target transpiration, indicating a more robust hydraulic architecture under contrasting recharge regimes. The observed xylem water potential per genotype for the mediterranean conditions follow experimental observations regarding the relative sensitivity of these genotypes to drought (Keller 2010): ‘SO4’ *>* ‘101-14’ *>*‘R110’.

### 4.4 Calibration under limited data

In the present study, the example calibration is constrained by the large age gap between the youngest (2-year-old) and oldest (48-year-old) vines. This limits our ability to assess whether the root system structure reaches a stable configuration by 48 years of age, or to resolve more gradual structural changes between 2 and 48 years that may arise from normal ontogenetic development or episodic stress. In addition, the mature and younger vines were grown at different sites and on different rootstocks, which may have contributed to differences in their RSAs independently of vine age. The present application should therefore be viewed primarily as an illustration of the modelling pipeline rather than as a comprehensive reconstruction of root-system development across the full lifespan of a vine. The model formulation draws partly on general patterns reported in the literature, while most of the empirical data used for calibration were obtained from the younger vines. Further data spanning a wider range of vine ages, ideally under comparable environmental and genetic conditions, would be needed to more rigorously constrain and evaluate age-dependent changes in root-system structure and turnover.

Several more general limitations regarding the data gathering pipelines for future studies should be acknowledged as well. Empirically, excavations of mature vines provide spatially truncated snapshots and are based on a limited number of individuals. Ring-based root ageing also requires caution: unlike stem wood, roots may exhibit eccentric growth, wedging or missing rings that can bias counts if not carefully evaluated (Cook and Kairiukstis 2013). In our system, clear annual boundaries and agreement between ring counts and known-age roots (1–4 years) support the interpretation of annual increments under strongly seasonal temperate conditions. However, ring expression may vary along the root axis and with soil depth, as well as with local carbon availability, and may differ across species and climates (De Micco et al. 2016).

### 4.5 Model simplifications

At the model level, structural root death is implemented as stochastic removal of entire subtrees once a lifetime threshold is exceeded. While this captures architectural consequences of turnover, it does not resolve underlying physiological mortality mechanisms such as hydraulic failure or carbon starvation (McDowell et al. 2008; McDowell et al. 2022; Sevanto et al. 2014). In addition, survival estimates derived from living-root age distributions assume approximate demographic stationarity; episodic disturbances (e.g. extreme drought or management changes) could transiently alter age structures and bias inferred turnover rates (Baden et al. 2021).

For the water flow simulation, the potential transpiration was prescribed as a constant daily demand, independent of vine age or short-term atmospheric variability. In reality, transpiration demand varies with canopy development, leaf area and meteorological conditions, and the investigated rootstocks differ substantially in aboveground vigour and associated transpirational demand (e.g. Fichtl et al. (2025)). The simulations therefore isolate belowground hydraulic capacity from dynamic shoot–root feedbacks and should be interpreted as a structural benchmark rather than a full representation of vine water use. Several simplifying assumptions deserve consideration. Vineyard soils are often constrained by shallow bedrock, compacted layers or abrupt textural transitions, which can limit effective rooting depth. Likewise, the imposed climatic scenarios represent idealised boundary conditions rather than realistic interannual variability. Importantly, these factors, including dynamic transpiration demand, genotype-specific canopy development, variable soil depth and more complex precipitation regimes, can be explicitly incorporated within the modelling framework for future *in silico* studies.

The hydraulic patterns should be interpreted in light of uncertainty in the *L*_pr_ parameters. They were obtained from the literature and set to the same value for all three rootstocks, based on the data of Cuneo et al. (2018), Gambetta et al. (2012), and Gambetta et al. (2013). In their study, Barrientos-Sanhueza et al. (2024) also looked at the *L*_pr_ of fine roots and found *L*_pr_ = 5 *×* 10*^−^*^8^ *m s^−^*^1^ *MPa^−^*^1^, about one order of magnitude lower than our value of 4 *×* 10*^−^*^7^ *m s^−^*^1^ *MPa^−^*^1^ (Gambetta et al. 2013). This discrepancy could be explained by the fact that, Gambetta et al. (2013) used root samples of 10 *mm* to 20 *mm* length while, in their study, Barrientos-Sanhueza et al. (2024) used roots ranging from 80 *mm* to 110 *mm*. Roots of this length were considered to be suberised roots in our simulation. Our *L*_pr_ of suberised roots (7 *×* 10*^−^*^8^ *m s^−^*^1^ *MPa^−^*^1^, Gambetta et al. (2012)) is indeed much closer to the ones observed by Barrientos-Sanhueza et al. (2024) for well-watered conditions. The high genetic and environmental plasticity of *L*_pr_ (Barrientos-Sanhueza et al. 2024) underlines the importance of reliable genotype conductivity data, potentially including a drought stress feedback loop.

Despite the simplified simulated processes and hydraulic parameter calibration, the simulations revealed consistent age- and genotype-dependent patterns. Accordingly, this study does not aim to predict vineyard water status under specific field conditions, but rather to demonstrate how hydraulically explicit perennial modelling can reveal genotype-specific vulnerabilities, buffering capacities and long-term hydraulic effects that remain inaccessible to short-term experiments. Taken together, our results challenge the notion of a permanent woody root scaffold in perennials and instead reveal perennial root systems as dynamic, age-structured populations undergoing continuous renewal over decades. By linking empirical demography to architecture and hydraulics, the framework offers a transferable approach for analysing long-term root system function and climate sensitivity across woody perennial crops and ecosystems.

### 4.6 Future directions for the calibration pipeline

A major bottleneck for perennial root modelling is not conceptual model structure, but the scarcity of empirical datasets capturing mature, field-grown root systems at sufficient depth and resolution. This limitation is widely recognised across root ecology, where belowground traits remain far less constrained than aboveground traits due to the destructive and labour-intensive nature of sampling (Tumber-Dávila et al. 2022). Progress will require coordinated acquisition of mature-system datasets across soils and climates, ideally integrating three-dimensional architecture with independent information on age, diameter and tissue status.

Although the data acquisition pipeline used for this study is intensive, it allowed for a precise calibration of the model’s structural parameters. One particularity of the grapevine production is that 90% of the world’s production is obtained from 10 different rootstocks (Bernardo et al. 2025; Keller 2010). Therefore, it remains possible to conduct an in-depth calibration (with root excavation) for those rootstocks. Using FSPM for modelling would then allow users to simulate the feedback of environmental conditions on growth (e.g., resource scarcity, soil structure). *in situ* measurement methods (e.g., isotopic measurements (Bernardo et al. 2025)) could then be used to re-calibrate pre-parameterised parameter sets in new study sites.

### 4.7 Future directions for the model development

On the modelling side, the present framework focuses on the effect of root system structure on plant water flow, without including feedback processes. This implementation provides a foundation for extensions that incorporate hydraulic–carbon feedbacks, variable soil depth constraints and dynamic atmospheric demand. Such developments would allow demographic turnover to respond plastically to stress rather than being prescribed solely by age. Other models used to simulate the growth of tree root systems give insights into processes which could be added to represent the root system structure and water transport of grapevines, such as the pruning of dead roots (Barczi et al. 2018), or the role of capacitance, cavitation and cuticular transpiration on the plant water balance (Cochard et al. 2021). The simulations in this study covered a 50-year range and had a resolution of 1 hour during the growing period to simulate the water flow. The field was represented implicitly by using periodic boundary conditions. The scenarios took less than one day to complete. This relatively low computation time enables the implementation of additional functional modules while keeping a simulation spanning several decades. Moreover, a multiscale pipeline was already implemented in CPlantBox to simulate the interaction between the carbon and water fluxes in the soil-rhizosphere plant continuum (Giraud et al. 2026). The same setup could be used for future grapevine simulations. Our simulations could also be used to inform higher scale models, for instance by providing the 1D-SUF, which is more closely related to the plant’s water uptake capacity than the root system structure itself.

## 5 Conclusion

Perennial root systems are fundamentally demographic systems, shaped by the continuous emergence, maturation and loss of absorptive and structural components over decades. By placing root age structure at the centre of model formulation and calibration, this study demonstrates that longterm root architecture and hydraulic function cannot be understood without explicitly accounting for age-dependent survival, growth regulation and secondary development.

Combining rare three-dimensional datasets from young and mature grapevines with dendrochrono-logical age reconstructions, we show that lignified structural roots do not form permanent scaffolds but instead undergo substantial turnover on decadal timescales. Embedding these empirical age distributions into a functional–structural modelling framework alters simulated trajectories of architecture, water uptake and hydraulic behaviour and reveals genotype-specific hydraulic strategies emerging over time.

Beyond grapevine, the conceptual advances presented here are broadly applicable to woody perennial species. Although further empirical data (e.g., to reach a higher temporal resolution) and model extensions (e.g., root-shoot coupling) are needed to resolve physiological mechanisms of root mortality and stress feedbacks, the framework provides a foundation for linking belowground demography with long-term plant function.

By moving perennial root modelling beyond static lifespan assumptions, this work establishes root age structure as a key organising principle for understanding resilience, hydraulic performance and climate sensitivity in woody perennial systems.

## Acknowledgements

We gratefully acknowledge the financial support of the German Federal Environmental Foundation (Deutsche Bundesstiftung Umwelt, DBU), and open access funding provided by Hochschule Geisenheim University. We further acknowledge funding by the German Research Foundation (DFG) under Germany’s Excellence Strategy, EXC-2070–390732324–PhenoRob. We thank Michael Bruch for his support during root system excavations. We are grateful to Jörg Kunz for advice on the dendrochronological analyses. We further thank David Eissenstat for helpful feedback on the interpretation of the dendrochronological results and woody root age determination. We thank the laboratory team of the Department of General and Organic Viticulture (HGU) for technical assistance. We further acknowledge Leonie Dries and Joschua Göttmann for their support with accompanying soil and plant analyses. We thank Jochen Eberbach from the Schloss Vollrads Wine Estate for granting access to the vineyard for excavation of mature root systems and for the excellent cooperation. Finally, we thank Katrin Kahlen and Dominik Schmidt for providing the digitisation equipment.

## Author Contributions

LF, MG, and MF conceived the study and developed the central research questions. LF, MG, and DL developed and implemented the modelling framework. LF and MG performed the analyses and simulations. LF, KS, and MF conducted the experimental work and data acquisition. LF and MG wrote the first draft of the manuscript. DL, AS, KS and MF contributed to data interpretation and critically revised the manuscript. AS and MF supervised the research. LF and MF acquired the funding. All authors contributed to the article and approved the submitted version.

## Data Availability

All simulations were performed using an extended version of the open-source framework CPlantBox (Giraud et al. 2023; Zhou et al. 2020). The code extensions described in this study were implemented in a dedicated branch of the official CPlantBox repository and are publicly available at https://github.com/Plant-Root-Soil-Interactions-Modelling/CPlantBox/tree/perennial2026 and https://github.com/Plant-Root-Soil-Interactions-Modelling/dumux-rosi/tree/perennial2026.

The grapevine 3D root architecture datasets used for model parameterisation and evaluation are openly available on Zenodo (Fichtl and Friedel 2025a,b) at: https://doi.org/10.5281/zenodo.15632062 and https://doi.org/10.5281/zenodo.17637194.

## Supplementary Information

The supplementary information includes the following materials:

### Supplementary Methods

**Methods S1:** Implementation of branching rules in CplantBox

**Methods S2:** Temperature-driven dormancy and BEDD implementation

**Methods S3:** Structural root survival, turnover and directional growth

**Methods S4:** Definition of segment age and implementation of secondary radial growth

**Methods S5:** Implementation of developmental status transitions

**Methods S6:** Hydraulic parameterisation

**Methods S7:** Estimation of maximum attainable structural root length

**Methods S8:** Age-dependent axial growth potential and thermal-time formulation

### Supplementary Figures

**Figure S1:** Order-specific root length distributions in young and mature grapevine root systems

**Figure S2:** Validation of simulated structural root survival probability distributions

**Figure S3:** Validation of simulated order-specific structural root numbers

### Supplementary Tables

**Table S1:** Order-specific Weibull parameters for structural root survival

**Table S2:** Axial and radial hydraulic parameters used in the model

**Table S3:** Order- and genotype-specific maximal root lengths and thermal sensitivity parameters

**Table S4:** Core and perennial CPlantBox model parameters

## Supplementary Methods

### Method S1: Implementation of branching rules in CPlantBox

Lateral initiation is evaluated at each potential branching node using rule-specific successor tables. The table entries comprise successorOT, which defines the organ type(s) of candidate laterals; successorST, which specifies the corresponding organ subtypes (here: structural order classes versus fine-root subtypes); successorNo, which sets the maximum number of laterals per insertion node; and successorP, which stores the probability vector over all candidate successor subtypes, with ∑*_i_ p_i_ ≤* 1.

At the moment of branching, CPlantBox calls the internal function getLateralType(). A uniform random number *d ∈* [0, 1] is drawn and compared against the cumulative sum of the entries in successorP. The first subtype for which the cumulative probability exceeds *d* is selected as the successor. Any residual probability mass 1*−∑_i_ p_i_* corresponds to the option of no lateral initiation at that node, in which case the function returns {-1,-1} and the parent segment remains unbranched.

To represent age-dependent shifts in allocation between structural and absorptive branching, we used successorP age to define discrete age thresholds (root system age in BEDD) for each branching rule (ruleId) at which the probability vector successorP is updated. Between thresholds, probability vectors are linearly interpolated as a function of root age using the internal interpolate P() routine. This allows a continuous transition from predominantly structural branching in young roots to increased fine-root production at intermediate ages, followed by a general decline in branching probability in older roots.

### Method S2: Temperature-driven dormancy and BEDD implementation

Dormancy and seasonal growth activity were implemented using a biologically effective degree day (BEDD) formulation commonly applied in viticulture (see section 2.3.10).

Daily values were summed over each calendar year to obtain an annual BEDD total,

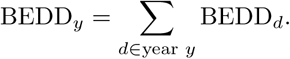

To ensure compatibility with the annual time-stepping scheme used in CPlantBox, annual BEDD totals were normalised by a reference seasonal heat sum,

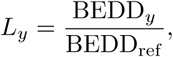

where BEDD_ref_ = 1225 BEDD corresponds to the long-term mean growing-season heat sum at the Geisenheim site. The resulting factor *L_y_* defines the effective number of growth iterations in year *y*.

Historical daily temperature records from the Geisenheim weather station were used to derive site-specific BEDD trajectories. Because the formulation is based on heat accumulation rather than fixed calendar dates, it can be readily applied to other climatic regions using locally available temperature data.

During BEDD-defined active periods, root elongation, branching and fine-root initiation were enabled. Outside this period, all growth processes were suppressed. At dormancy onset, all newly formed unsuberised fine roots were removed, while structural roots persisted and were subject only to their age-dependent survival functions.

### Method S3: Structural root survival, turnover, and directional growth

#### Weibull-based survival model for structural roots

Age-dependent survival of long-lived structural roots was implemented as a stochastic lifetime process based on order-specific Weibull distributions. For each newly created structural root of order *o*, an individual lifetime *rlt*_winter_ (in BEDD) was drawn at the time of root initiation as

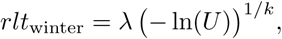

where *U ∼* Uniform(0, 1), and *λ* (in BEDD) and *k* (*−*) denote the scale and shape parameters of the Weibull distribution for root order *o*. Parameter values were derived from Kaplan–Meier survival curves fitted to dendrochronologically reconstructed basal root ages.

This formulation assigns each structural root a fixed maximum lifespan at creation, which is independent of subsequent environmental conditions or local root density. Survival is evaluated annually at the onset of winter dormancy. If the chronological age of a root exceeds its assigned lifetime, the root is removed from the system.

The results of the calibration, based on the observed data (grey ribbons in Figure S2) can be seen in Table S1. Differences in parameters between root types were evaluated using a likelihood-ratio test (LRT). The full model allowed each root type to have its own *k* and *λ*. This model was tested against another simpler model for which *k* and *λ* were both shared across root types (null hypothesis: the survival curves are identical). We set the significance level *α* to 0.05. The null hypothesis could always be rejected, especially for “Main” against the other two root groups (*p <* 1*e^−^*^8^). The difference between “Sub” and “Subsub” was significant but much nearer the threshold (*p* = 0.026).

The *λ* parameter defines the age at which a fraction of *e^−^*^1^ = 0.37 of the roots remains alive, while *k* defines the shape of the curve around this value (how mortality varies with age). We observe that the *k* parameter is similar between the root orders while the *λ* changes more strongly. This indicates that root mortality follows similar patterns between root order, though high-order roots have shorter life spans.

#### Recursive removal of structural subtrees

When a structural root exceeds its assigned Weibull lifetime, the root and its entire downstream subtree are removed recursively. This includes all descendant structural and fine roots emerging from that root. This design choice reflects the biological reality that death of a woody transport axis typically renders all dependent distal tissues non-functional. Partial segment-level mortality was not implemented, as empirical data do not resolve such fine-scale persistence in mature woody roots.

In addition to age-dependent mortality, structural roots that fail to reach a minimum lignified length before the onset of dormancy are removed. This threshold is controlled by the parameter Lmax unsuberised and ensures that only sufficiently developed roots persist across winter.

#### Emergent demographic structure without explicit balancing

Structural root recruitment is governed exclusively by local branching rules defined in the successor tables (Methods S1). No explicit demographic balancing, target root number, or age-distribution forcing is applied. Instead, population-level age distributions emerge from the interaction of stochastic lifetime assignment, recursive mortality, and age-dependent branching probabilities. As a consequence, the simulated perennial root system approaches a dynamic steady state in which continuous recruitment of new structural roots is balanced by stochastic loss of older ones, without requiring any global constraints on root abundance or age structure.

#### Directional growth of structural roots: probabilistic tropism

Directional growth of structural roots was controlled using a stochastic mixture tropism implemented as a custom tropism type (tt shift), following the approach introduced for grapevine root systems by Fichtl et al. (2024). This formulation combines two existing tropism functions—gravitropism and plagiotropism—via probabilistic switching.

For each newly created root segment, a uniform random number *U ∼* Uniform(0, 1) is drawn and compared to the cumulative tropism weights tropismW1 and tropismW2. Depending on this draw, either the gravitropic or the plagiotropic objective function is evaluated and minimised to determine the growth direction of the segment. Thus, each segment follows a single tropistic objective, but the population of segments exhibits a mixed directional response.

Tropism weights were assigned in an order- and genotype-specific manner. Primary axes (“main” roots) were parameterised with mixed gravitropic and plagiotropic tendencies (e.g. 40/60 weighting), whereas higher-order structural roots were biased towards gravitropic growth. This formulation reproduces the observed coexistence of deep, vertically oriented transport roots and laterally spreading structural branches in grapevine root systems. The mixture tropism is fully compatible with the existing CPlantBox tropism infrastructure and introduces stochastic architectural variability among replicate simulations even under identical parameter sets.

### Method S4: Definition of segment age and implementation of secondary radial growth

Secondary radial growth in CPlantBox is implemented at the level of individual root segments and depends on segment age rather than on global simulation time. For each segment, age is derived from the chronological age of the parent root and the creation times of the segment-defining nodes. Let *a*_root_ (in BEDD) denote the age of the root (time since root initiation), *t*_node,0_ (BEDD) the creation time of the basal node of the root, and *t*_node,seg_ (BEDD) the creation time of the node terminating the segment. Segment age is then computed as

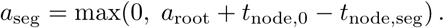

This definition ensures that segments formed earlier along a root axis (i.e. basal segments) accumulate more secondary growth than younger, more distal segments. Negative values, which may arise for newly created apical segments, are truncated at zero.

Secondary radial growth is specified as a linear function of segment age (see Eqn. 4). Parameters were assigned independently for each structural root order based on empirical age–diameter relationships derived from dendrochronological analyses.

Segment radii are evaluated dynamically whenever required by geometric or hydraulic routines and are not stored as explicit state variables. Consequently, secondary growth terminates automatically when a root exceeds its stochastic lifetime (drawn from the order-specific Weibull distribution) and is removed from the simulation, without the need for additional bookkeeping.

### Method S5: Implementation of developmental status transitions

Developmental status transitions were implemented using a three-state scheme representing key anatomical zones along grapevine roots, following Gambetta et al. (2013). The states correspond to (i) unsuberised tissue in the meristematic and elongation zone, (ii) suberised tissue in the maturation zone, and (iii) lignified tissue undergoing secondary growth.

Rather than tracking anatomical markers explicitly, developmental status is determined by cumulative length of root tissue produced distal to a given segment, which provides a robust and computationally efficient proxy for axial tissue maturation. Two fixed length thresholds were defined:

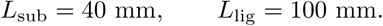

For a root segment *i* with the length between its apical node and the tip of the root *L_i_*(*t*), developmental status is assigned as

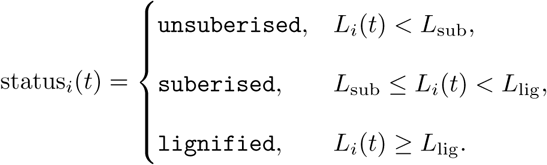

Unsuberised segments are characterised by high radial hydraulic conductivity and are eligible for fine-root turnover at dormancy onset. Suberised segments exhibit reduced radial permeability due to apoplastic barrier formation. Lignified segments are eligible for secondary radial growth and represent long-lived woody tissue. Status transitions are irreversible and evaluated dynamically as roots elongate.

### Method S6: Hydraulic parameterisation

Root hydraulic properties were implemented by assigning axial conductances (*K*_h_, *m*^4^ *s^−^*^1^ *MPa^−^*^1^) and radial conductivities (*L*_pr_, *m s^−^*^1^ *MPa^−^*^1^) to each root segment based on its simulated diameter, age, and developmental status.

#### Axial hydraulic conductances

*K*_h_ of structural roots was computed as a power-law function of external root diameter *D*,

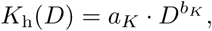

where *D* is the simulated outer root diameter (m), and *a_K_*(*m*^3^ *s^−^*^1^ *MPa^−^*^1^) and *b_K_* (*−*) are genotype-specific parameters. These relationships were derived by Fichtl et al. (2024) from anatomical analyses of grapevine rootstocks, combining measurements of xylem vessel number and size with the Hagen–Poiseuille equation adapted for elliptical conduits. Separate fits were obtained for each rootstock genotype using nonlinear least squares and showed high predictive accuracy across a wide diameter range. In the present model, *K*_h_ was evaluated dynamically for each segment based on its age-dependent diameter.

#### Radial hydraulic conductivity

*L*_pr_ was assigned based on the developmental status of each root segment (unsuberised, suberised, lignified; see Methods S5). Because rootstock-specific *L*_pr_ measurements are not available, representative literature values for grapevine roots were used. These values capture the strong reduction in radial permeability associated with suberisation and lignification and were applied uniformly across genotypes.

#### Hydraulic coupling

*K*_h_ and *L*_pr_ were integrated into the CPlantBox hydraulic solver to compute segment-specific water uptake, flow redistribution within the root system, and whole-root-system conductance (*K*_rs_, *m*^4^ *s^−^*^1^ *MPa^−^*^1^). *K*_h_ increases with segment age due to secondary thickening, whereas *L*_pr_ decreases as roots transition from unsuberised to lignified tissue. This formulation links root architectural development, anatomical maturation, and hydraulic function in a mechanistic and internally consistent manner.

### Method S7: Estimation of maximum attainable structural root length

Maximum attainable lengths of structural roots were estimated from the young vine excavation dataset, which provides time-resolved three-dimensional root system architectures during the first two years after planting. Because full root system extents cannot be observed directly over multi-decadal timescales, we inferred characteristic upper length bounds from early elongation trajectories.

For each rootstock genotype and structural branching order, root lengths were extracted from the longest observed roots at each excavation time point. Root age was expressed as time since the appearance of the respective order. For “main” and “sub” roots, age was defined as time since planting, whereas for “subsub” roots (orders 4–6), which emerged only from the second growing season onward, age was defined relative to their first observed occurrence to avoid negative ages.

For each rootstock *×* order combination, an asymptotic growth function (See Eqn. 2) was fitted using nonlinear least squares. Fits were constrained such that *L*_max_ (*cm*) was at least as large as the maximum observed length *L*_max*,o*_ (*cm*) and *β >* 0. Combinations with fewer than two distinct age levels were excluded from fitting.

The fitted asymptote was interpreted as the maximum attainable length for the corresponding rootstock and structural order. For all fitted combinations, predicted lengths approached the asymptote within approximately 25 years, indicating that the inferred maximum lengths represent decadal-scale elongation limits rather than short-term artefacts. These *L*_max_ values were used as order-specific parameters in the age-dependent growth potential formulation.

### Method S8: Age-dependent axial growth potential and thermal-time formulation

To derive a climate-driven representation of age-dependent root elongation, we linked empirically inferred growth damping to cumulative biologically effective degree days (BEDD). This approach allows age effects to be expressed implicitly through thermal-time saturation rather than explicit age tracking during simulation.

For each excavated structural root of the mature grapevine dataset, we estimated a hypothetical total root length, denoted 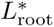. This quantity represents the length a root would have reached in the absence of age-dependent growth damping and is used here solely as an intermediate variable to infer growth–climate relationships. Importantly, 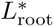 is distinct from the order-specific maximum lengths *l*_max_ used as model input, which were independently derived from young-vine architectural data.

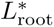 was estimated by inverting the empirically derived cumulative damping function *χ*(*a*) along all segments of a root. For a segment spanning ages *a*_apical_ to *a*_basal_, the expected segment length *ℓ* is given by

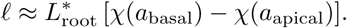

Individual values of 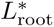 were obtained either by robust regression of *ℓ* on Δ*χ* across segments or by computing the root-wise median of *ℓ/*Δ*χ*. Aggregation across roots yielded order-specific distributions of 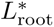.

For each sampled root, cumulative BEDD experienced over its lifetime was reconstructed by summing annual BEDD values backwards from the sampling year according to basal root age (see Dormancy formulation). This yielded a root-specific thermal exposure BEDD_tot_.

Estimated total root length was then related to BEDD_tot_ using a saturating exponential model,

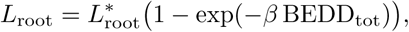

where *β* is an order-specific thermal sensitivity parameter estimated by nonlinear least squares. In this formulation, the exponential complement exp(*−β* BEDD) represents the remaining growth potential in thermal-time space.

By construction, the fitted BEDD-based saturation curves reproduce the empirically derived age-dependent relative growth potential (RGP) under the observed temperature history, thereby embedding age-dependent growth decline into a climate-driven formulation.

In the CPlantBox simulations, elongation of structural roots is therefore controlled solely by accumulated BEDD, *L*_max_, and *β*. At each annual time step, root length is incremented according to the additional BEDD accumulated during that year. This formulation avoids explicit evaluation of age-dependent growth damping during runtime while preserving the empirically constrained decline in elongation capacity with increasing root age.

## Supplementary Figures

**Figure S1:**
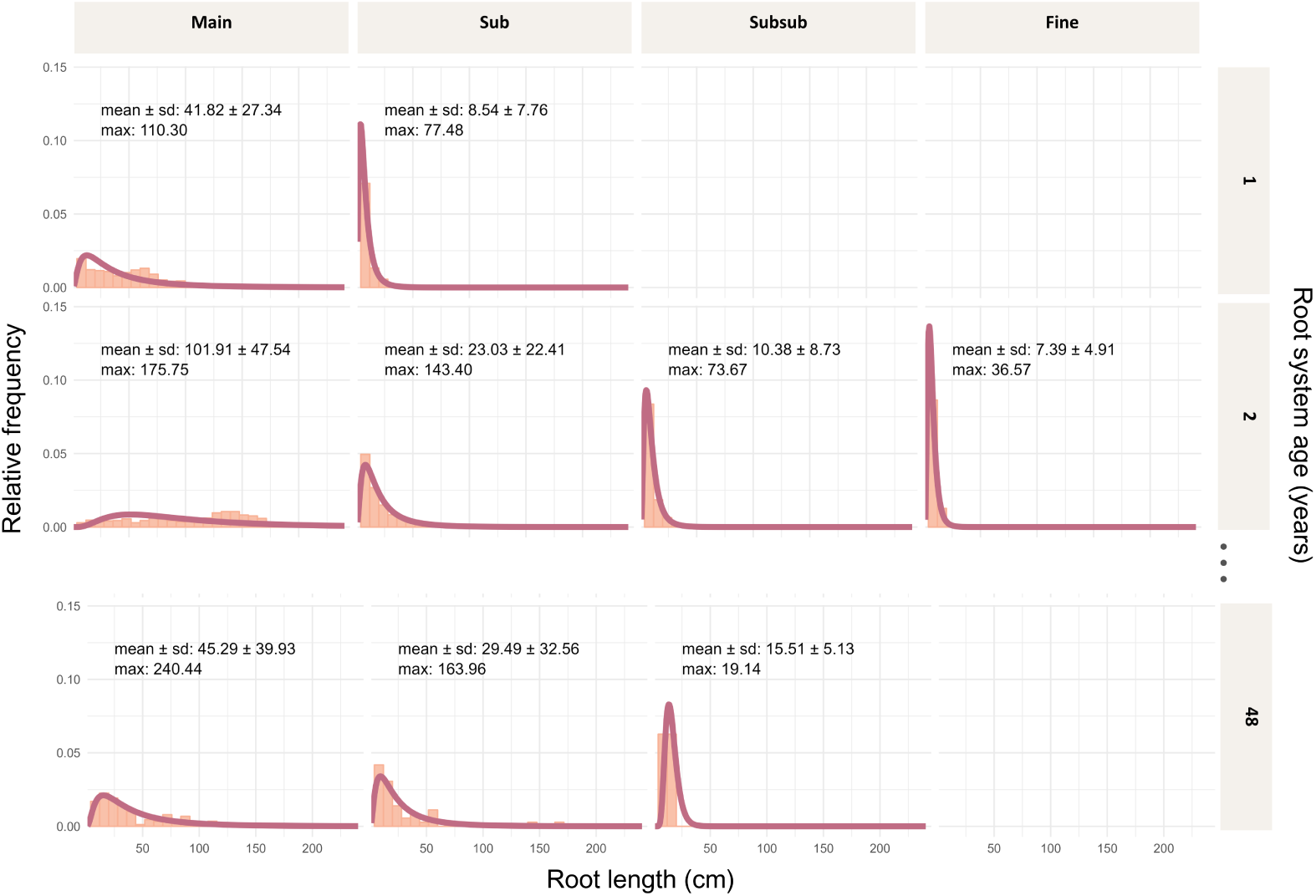
Order-specific root length distributions from young (1–2 years) and 48-year-old grapevine root systems. Note that root lengths in the mature vines are truncated by the excavation volume and therefore represent partial system lengths.

**Figure S2:**
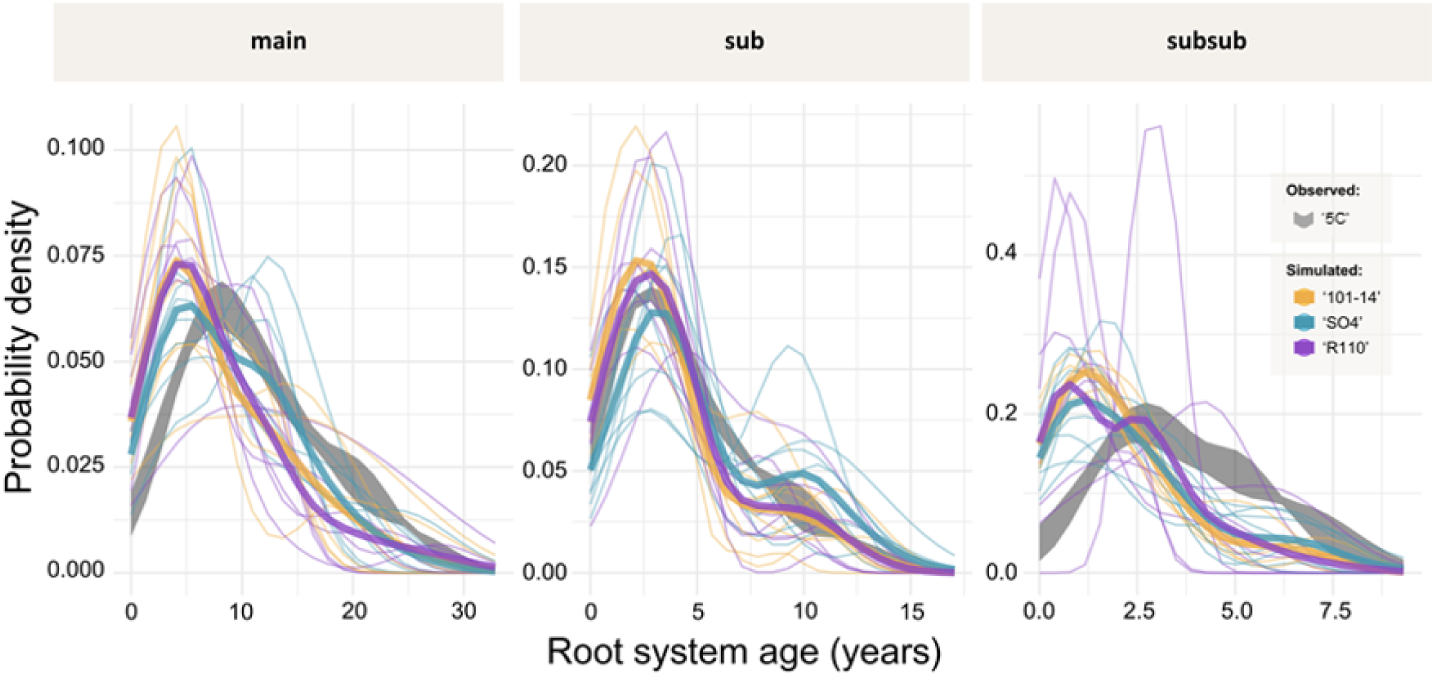
Validation of simulated structural root survival probability distributions with Kernel density estimates of survival probability for “main”, “sub” and “subsub” roots (three rootstocks: ‘101-14’,‘SO4’, and ‘R110’). Grey ribbons show observed distributions (mean *±* SD) derived from dendrochronological data (rootstock ‘5C’). Coloured thin lines represent individual simulation runs (*n* = 8 per rootstock), and thick coloured lines indicate genotype-specific means.

**Figure S3:**
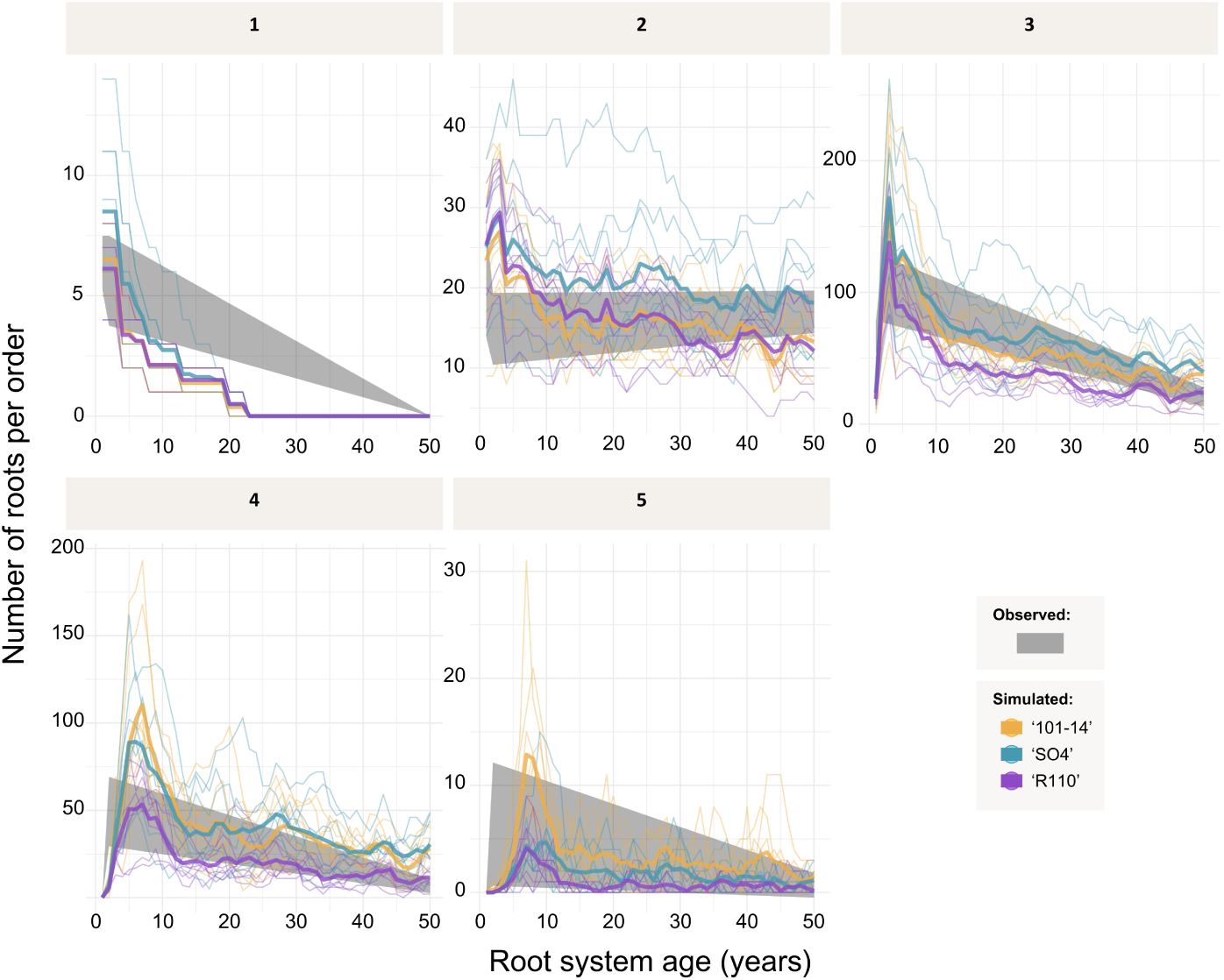
Validation of simulated order-specific structural root numbers. Grey ribbons indicate observed root numbers derived from young (three rootstocks: ‘101-14’,‘SO4’, and ‘R110’) and mature 3D RSA (rootstock ‘5C’). Coloured thin lines show individual simulation runs (*n* = 8 per genotype), and thick coloured lines represent genotype-specific mean trajectories. In mature vines, first-order roots (adventitious roots present at planting) were no longer observed; consequently, newly emerging permanent structural roots were classified as second-order roots.

## Supplementary Tables

**Table S1:** Order-specific Weibull parameters describing survival of structural grapevine roots. Parameters were estimated from Kaplan–Meier survival curves shown in Fig. S2. Distinct parameter values reflect strong contrasts in longevity between “main”, “sub” and “subsub” roots.

| Root order | Shape $k$ | Scale $\lambda$ (years) | Max. age $A_{\max}$ (years) |
| --- | --- | --- | --- |
| “Main” | 1.88 | 7.53 | 23 |
| “Sub” | 1.42 | 3.54 | 14 |
| “Subsub” | 2.17 | 4.55 | 8 |

**Table S2:** Axial and radial hydraulic parameters used in the perennial root architecture model. Diameter values refer to the external root diameter (m).

| Parameter | Function / Value | Unit | Reference |
| --- | --- | --- | --- |
| **Axial hydraulic conductance ($K_h$)** |  |  |  |
| ‘101-14’ | $K = 0.04749 \cdot D^{2.06437}$ | $\text{m}^4 \text{s}^{-1} \text{MPa}^{-1}$ | Fichtl et al. (2024) |
| ‘SO4’ | $K = 0.1539 \cdot D^{2.2410}$ | $\text{m}^4 \text{s}^{-1} \text{MPa}^{-1}$ | Fichtl et al. (2024) |
| ‘R110’ | $K = 0.02622 \cdot D^{1.98847}$ | $\text{m}^4 \text{s}^{-1} \text{MPa}^{-1}$ | Fichtl et al. (2024) |
| **Radial hydraulic conductivity ($L_{\text{pr}}$)** |  |  |  |
| Lignified root | $4.0 \times 10^{-8}$ | $\text{m s}^{-1} \text{MPa}^{-1}$ | Cuneo et al. (2018) |
| Suberised root | $7.0 \times 10^{-8}$ | $\text{m s}^{-1} \text{MPa}^{-1}$ | Gambetta et al. (2012) |
| Unsuberised root | $4.0 \times 10^{-7}$ | $\text{m s}^{-1} \text{MPa}^{-1}$ | Gambetta et al. (2013) |

**Table S3:** Order- and genotype-specific maximal root lengths (*L*_max_) derived from young-vine growth trajectories, and thermal sensitivity parameters (*β*) governing the exponential damping of elongation with accumulated biologically effective degree days (BEDD). *L*_max_ is genotype-specific, whereas *β* varies by structural order.

| Root order | Rootstock | $L_{\max}$ (cm) | $\beta$ (BEDD) |
| --- | --- | --- | --- |
| “Main” | ‘R110’ | 790 | 0.10 |
|  | ‘101-14’ | 630 | 0.07 |
|  | ‘SO4’ | 640 | 0.09 |
| “Sub” | ‘R110’ | 120 | 0.02 |
|  | ‘101-14’ | 100 | 0.02 |
|  | ‘SO4’ | 90 | 0.02 |
| “Subsub” | ‘R110’ | 60 | 0.02 |
|  | ‘101-14’ | 50 | 0.02 |
|  | ‘SO4’ | 50 | 0.02 |

**Table S4:**
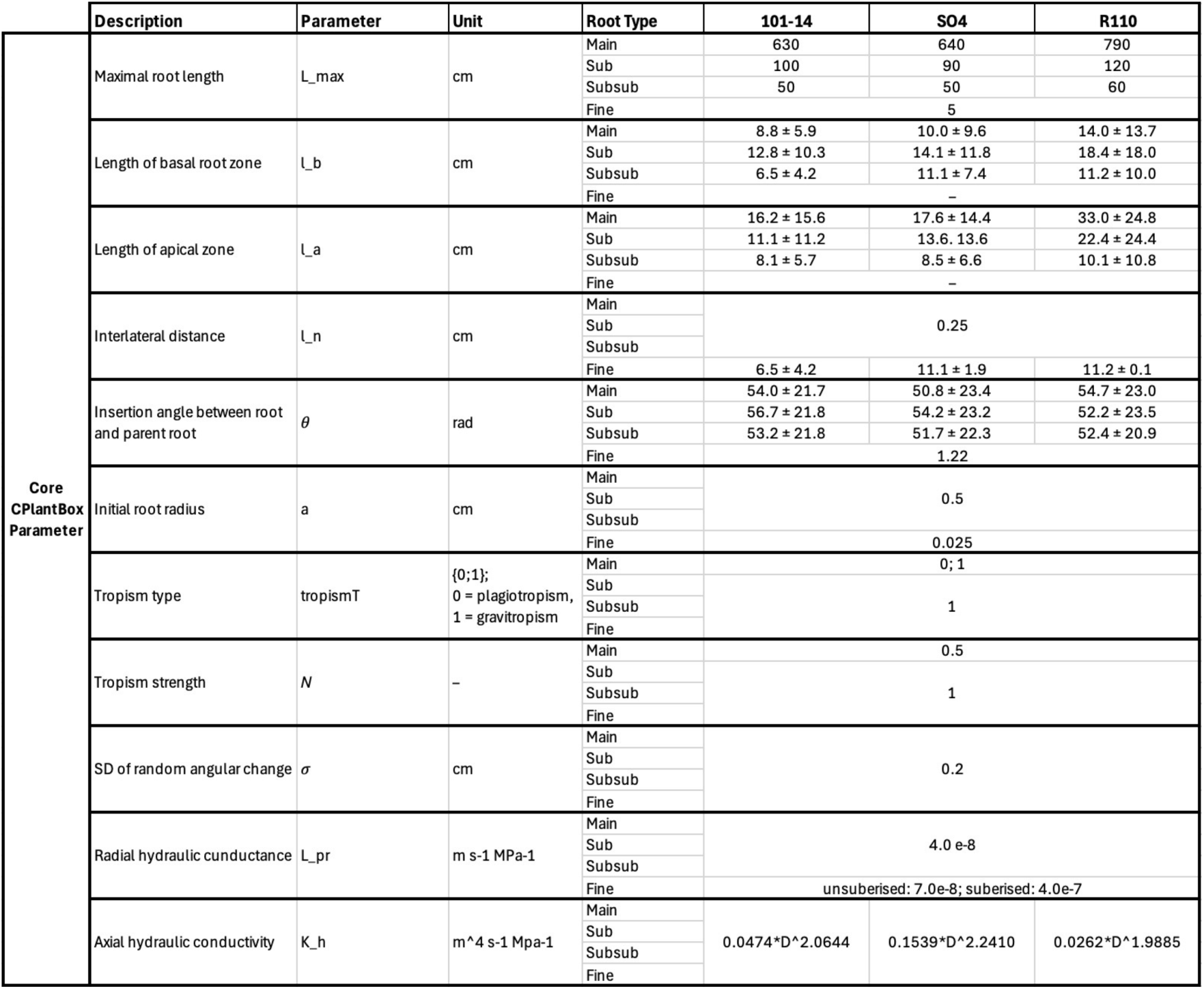
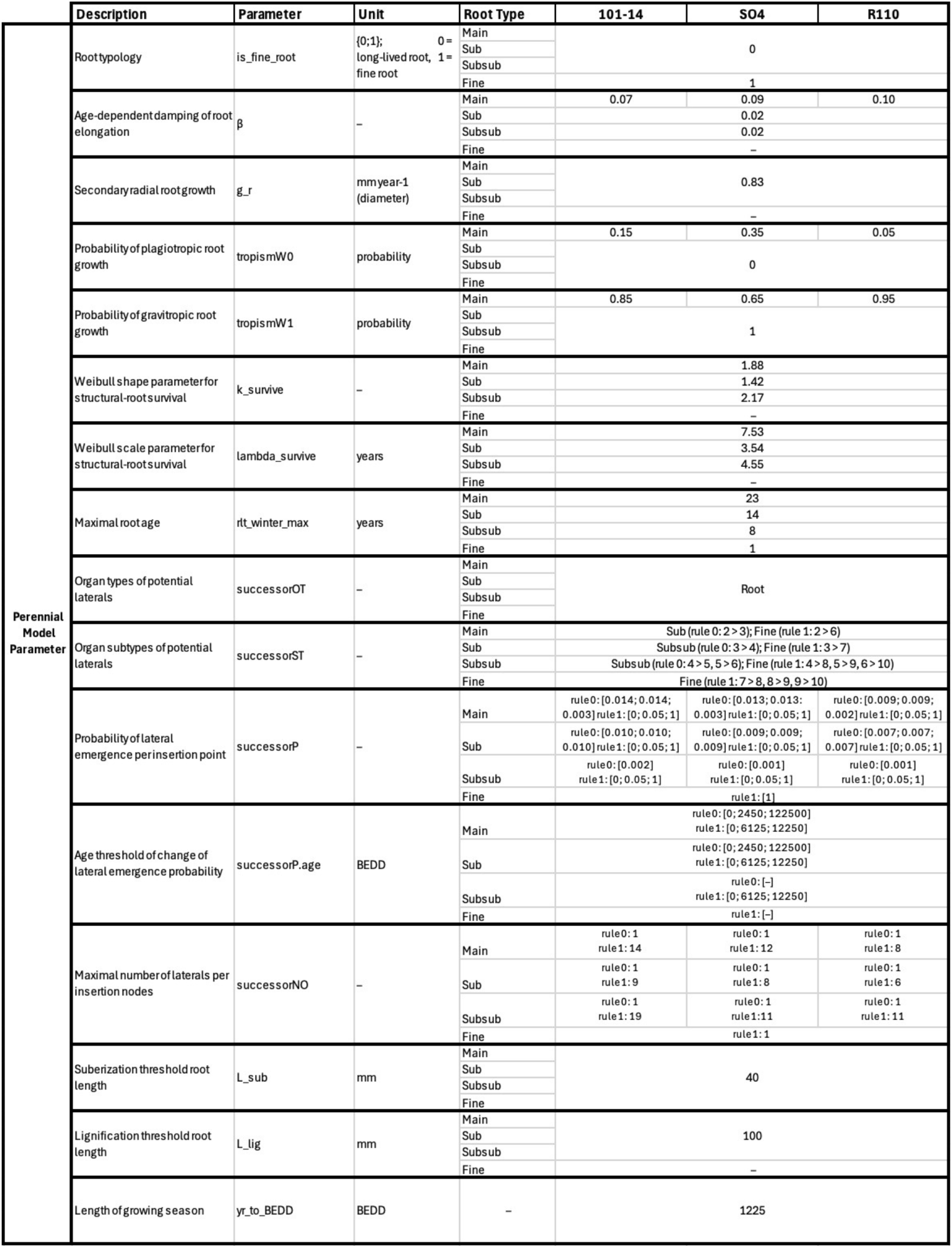
Core CPlantBox parameters used for perennial root system architecture (RSA) simulations of three grapevine rootstock genotypes. Perennial CPlantBox parameters used for perennial root system architecture (RSA) simulations of three grapevine rootstock genotypes.

## References

Ahrens, J., B. Geveci, C. Law, C. Hansen, and C. Johnson (2005). ParaView: An end-user tool for large-data visualization. In: The visualization handbook 717, pp. 50038–1.

Anderson, L. J., L. Comas, A. Lakso, and D. M. Eissenstat (2003). Multiple risk factors in root survivorship: a 4-year study in Concord grape. In: New phytologist, pp. 489–501.

Baden, H. M., D. Roach, F. H. Schweingruber, K. Reitzel, K. Lundgreen, and J. P. Dahlgren (2021). The effects of age on the demography of a perennial plant depend on interactions with size and environment. In: Journal of Ecology 109.2, pp. 1068–1077.

Barczi, J.-F., H. Rey, S. Griffon, and C. Jourdan (Apr. 2018). DigR: a generic model and its open source simulation software to mimic three-dimensional root-system architecture diversity. In: Annals of Botany 121.5, pp. 1089–1104. issn: 0305-7364. doi: 10.1093/aob/mcy018. eprint: https://academic.oup.com/aob/article-pdf/121/5/1089/28751137/mcy018.pdf. url: 10.1093/aob/mcy018.

Barrientos-Sanhueza, C., A. Zurita-Silva, T. Knipfer, A. McElrone, and I. Cuneo (Aug. 2024). Unique root hydraulic and mechanical properties support the resilience of grapevines adapted to the Atacama Desert. In: Plant, Cell & Environment 47, pp. 5126–5139. doi: 10.1111/pce.15085.

Bates, T. R., R. M. Dunst, and P. Joy (Jan. 2002). Seasonal Dry Matter, Starch, and Nutrient Distribution in ‘Concord’ Grapevine Roots. In: HortScience 37.2, pp. 313–316. issn: 0018-5345. doi: 10.21273/HORTSCI.37.2.313. url: 10.21273/HORTSCI.37.2.313.

Bauerle, T. L., D. R. Smart, W. L. Bauerle, C. Stockert, and D. M. Eissenstat (2008). Root foraging in response to heterogeneous soil moisture in two grapevines that differ in potential growth rate. In: New Phytologist 179.3, pp. 857–866.

Bernardo, S., E. Marguerit, N. Ollat, G. A. Gambetta, C. Saint Cast, and M. de Miguel (Aug. 2025). Root system ideotypes: what is the potential for breeding drought-tolerant grapevine rootstocks? In: Journal of Experimental Botany 76.11, pp. 2970–2984. issn: 0022-0957. doi: 10.1093/jxb/eraf006. eprint: https://academic.oup.com/jxb/article-pdf/76/11/2970/61398043/eraf006.pdf. url: 10.1093/jxb/eraf006.

Cochard, H., F. Pimont, J. Ruffault, and N. Martin-StPaul (June 2021). SurEau: a mechanistic model of plant water relations under extreme drought. In: Annals of Forest Science 78. doi: 10.1007/s13595-021-01067-y.

Collet, C., M. Löf, and L. Pagès (2006). Root system development of oak seedlings analysed using an architectural model. Effects of competition with grass. In: Plant and Soil 279.1, pp. 367–383.

Comas, L. H., T. Bauerle, and D. Eissenstat (2010). Biological and environmental factors controlling root dynamics and function: effects of root ageing and soil moisture. In: Australian Journal of Grape and Wine Research 16, pp. 131–137.

Comas, L. H., D. M. Eissenstat, and A. N. Lakso (2000). Assessing root death and root system dynamics in a study of grape canopy pruning. In: The New Phytologist 147.1, pp. 171–178.

Cook, E. R. and L. A. Kairiukstis (2013). Methods of dendrochronology: applications in the environmental sciences. Springer Science & Business Media.

Couvreur, V., J. Vanderborght, and M. Javaux (2012). A simple three-dimensional macroscopic root water uptake model based on the hydraulic architecture approach. In: Hydrology and earth system sciences 16.8, pp. 2957–2971.

Cuneo, I. F., T. Knipfer, P. Mandal, C. R. Brodersen, and A. J. McElrone (2018). Water uptake can occur through woody portions of roots and facilitates localized embolism repair in grapevine. In: New Phytologist 218.2, pp. 506–516.

de Dorlodot, S., B. Forster, L. Pagès, A. Price, R. Tuberosa, and X. Draye (2007). Root system architecture: opportunities and constraints for genetic improvement of crops. In: Trends in plant science 12.10, pp. 474–481.

De Micco, V., F. Campelo, M. De Luis, A. Bräuning, M. Grabner, G. Battipaglia, and P. Cherubini (2016). Intra-annual density fluctuations in tree rings: how, when, where, and why? In: Iawa Journal 37.2, pp. 232–259.

Delrot, S., J. Grimplet, P. Carbonell-Bejerano, A. Schwandner, P.-F. Bert, L. Bavaresco, L. D. Costa, G. Di Gaspero, E. Duchêne, L. Hausmann, M. Malnoy, M. Morgante, N. Ollat, M. Pecile, and S. Vezzulli (2020). “Genetic and Genomic Approaches for Adaptation of Grapevine to Climate Change”. In: Genomic Designing of Climate-Smart Fruit Crops. Ed. by C. Kole. Cham: Springer International Publishing, pp. 157–270. isbn: 978-3-319-97946-5. doi: 10.1007/978-3-319-97946-5_7. url: 10.1007/978-3-319-97946-5_7.

Diggle, A. (1988). ROOTMAP—a model in three-dimensional coordinates of the growth and structure of fibrous root systems. In: Plant and soil 105.2, pp. 169–178.

Dunbabin, V. M., J. A. Postma, A. Schnepf, L. Pagès, M. Javaux, L. Wu, D. Leitner, Y. L. Chen, Z. Rengel, and A. J. Diggle (2013). Modelling root–soil interactions using three–dimensional models of root growth, architecture and function. In: Plant and Soil 372.1, pp. 93–124. issn: 1573-5036. doi: 10.1007/s11104-013-1769-y. url: 10.1007/s11104-013-1769-y.

Dupuy, L., P. J. Gregory, and A. G. Bengough (2010). Root growth models: towards a new generation of continuous approaches. In: Journal of experimental botany 61.8, pp. 2131–2143.

Elzhov, T. V., K. M. Mullen, A.-N. Spiess, and B. Bolker (2023). minpack.lm: R Interface to the Levenberg-Marquardt Nonlinear Least-Squares Algorithm Found in MINPACK, Plus Support for Bounds. R package version 1.2-4. url: https://CRAN.R-project.org/package=minpack.lm.

Fichtl, L. and M. Friedel (2025a). Grapevine3D: A Root System Architecture Dataset of 48-Year-Old Field-Grown Grapevines. Version 1.0. Zenodo. url: 10.5281/zenodo.17637194.

Fichtl, L. and M. Friedel (2025b). Grapevine3D: A Whole-Plant Architecture Dataset of Grapevine. Version 1.0. Zenodo. url: 10.5281/zenodo.15632062.

Fichtl, L., M. Hofmann, K. Kahlen, K. P. Voss-Fels, C. S. Cast, N. Ollat, P. Vivin, S. Loose, M. Nsibi, J. Schmid, et al. (2023). Towards grapevine root architectural models to adapt viticulture to drought. In: Frontiers in Plant Science 14, p. 1162506.

Fichtl, L., D. Leitner, A. Schnepf, D. Schmidt, K. Kahlen, and M. Friedel (2024). A field-to-parameter pipeline for analyzing and simulating root system architecture of woody perennials: application to grapevine rootstocks. In: Plant Phenomics 6, p. 0280.

Fichtl, L., K. Steng, A. Schnepf, and M. Friedel (2025). Rootstock genotype shapes whole-plant 3-D architecture and biomass allocation in field-grown grapevines. In: Annals of Botany 136.7, pp. 1613–1629.

Freschet, G. T., C. Roumet, L. H. Comas, M. Weemstra, A. G. Bengough, B. Rewald, R. D. Bardgett, G. B. De Deyn, D. Johnson, J. Klimešová, et al. (2021). Root traits as drivers of plant and ecosystem functioning: current understanding, pitfalls and future research needs. In: New Phytologist 232.3, pp. 1123–1158.

Friedel, M., M. Hendgen, M. Stoll, and O. Löhnertz (2020). Performance of reflectance indices and of a handheld device for estimating in-field the nitrogen status of grapevine leaves. In: Australian Journal of Grape and Wine Research 26.2, pp. 110–120.

Gambetta, G. A., C. M. Manuck, S. T. Drucker, T. Shaghasi, K. Fort, M. A. Matthews, M. A. Walker, and A. J. McElrone (Nov. 2012). The relationship between root hydraulics and scion vigour across Vitis rootstocks: what role do root aquaporins play? In: Journal of Experimental Botany 63.18, pp. 6445–6455. issn: 0022-0957. doi: 10.1093/jxb/ers312. eprint: https://academic.oup.com/jxb/article-pdf/63/18/6445/16932689/ers312.pdf. url: 10.1093/jxb/ers312.

Gambetta, G. A., J. Fei, T. L. Rost, T. Knipfer, M. A. Matthews, K. A. Shackel, M. A. Walker, and A. J. McElrone (2013). Water uptake along the length of grapevine fine roots: developmental anatomy, tissue-specific aquaporin expression, and pathways of water transport. In: Plant Physiology 163.3, pp. 1254–1265.

Giraud, M., A. K. Sırcan, T. Streck, D. Leitner, G. Lobet, H. Pagel, and A. Schnepf (2026). In silico analysis of carbon and water dynamics in the rhizosphere under drought conditions. In: SOIL 12.1, pp. 371–419. doi: 10.5194/soil-12-371-2026. url: https://soil.copernicus.org/articles/12/371/2026/.

Giraud, M., S. L. Gall, M. Harings, M. Javaux, D. Leitner, F. Meunier, Y. Rothfuss, D. van Duss-choten, J. Vanderborght, H. Vereecken, et al. (2023). CPlantBox: a fully coupled modelling platform for the water and carbon fluxes in the soil–plant–atmosphere continuum. In: in silico Plants 5.2, diad009.

Gladstones, J. (2021). Viticulture and Environment: A Study of the Effects of Environment on Grapegrowing and Wine Qualities, with Emphasis on Present and Future Areas for Growing Winegrapes. 2nd. Australia: Trivinum Press Pty Limited.

Jourdan, C. and H. Rey (1997). Modelling and simulation of the architecture and development of the oil-palm (Elaeis guineensis Jacq.) root system. In: Plant and soil 190.2, pp. 217–233.

Kaplan, E. L. and P. Meier (1958). Nonparametric estimation from incomplete observations. In: Journal of the American statistical association 53.282, pp. 457–481.

Kassambara, A., M. Kosinski, and P. Biecek (2025). survminer: Drawing Survival Curves using ‘ggplot2’. R package version 0.5.1. url: https://rpkgs.datanovia.com/survminer/index.html.

Keller, M. (2010). Chapter 1 - Botany and Anatomy. In: The Science of Grapevines. San Diego: Academic Press, pp. 1–47. isbn: 978-0-12-374881-2. doi: 10.1016/B978-0-12-374881-2.00001-5. url: https://www.sciencedirect.com/science/article/pii/B9780123748812000015.

Lanari, V., L. Pallotti, T. Lattanzi, and O. Silvestroni (2025). Grapevine Root Distribution and Density in Deep Soil Layers Under Different Soil Management Practices. In: Plants 14.12, p. 1823.

Leitner, D., S. Klepsch, G. Bodner, and A. Schnepf (2010). A dynamic root system growth model based on L-Systems: Tropisms and coupling to nutrient uptake from soil. In: Plant and soil 332.1, pp. 177–192.

Linsenmeier, A. (Jan. 2007). Einfluss der Stickstoffversorgung der Rebe (Vitis vinifera L. cv. Riesling) auf den untypischen Alterungston. In: Geisenheim: Forschungsanstalt Geisenheim 2007 (Geisenheimer Berichte; 60).

Linsenmeier, A. W., U. Loos, and O. Löhnertz (2008). Must composition and nitrogen uptake in a long-term trial as affected by timing of nitrogen fertilization in a cool-climate Riesling vineyard. In: American Journal of Enology and Viticulture 59.3, pp. 255–264.

Lobet, G., M. P. Pound, J. Diener, C. Pradal, X. Draye, C. Godin, M. Javaux, D. Leitner, F. Meunier, P. Nacry, et al. (2015). Root system markup language: toward a unified root architecture description language. In: Plant physiology 167.3, pp. 617–627.

Lynch, J. (1995). Root architecture and plant productivity. In: Plant physiology 109.1, p. 7.

Lynch, J. P., K. L. Nielsen, R. D. Davis, and A. G. Jablokow (1997). SimRoot: modelling and visualization of root systems. In: Plant and Soil 188.1, pp. 139–151.

Mahmud, K. P., S. K. Field, S. Y. Rogiers, S. Nielsen, Y. Guisard, and B. P. Holzapfel (2023). Rootstocks alter the seasonal dynamics and vertical distribution of new root growth of Vitis vinifera cv. Shiraz grapevines. In: Agronomy 13.9, p. 2355.

Mapfumo, E., D. Aspinall, and T. W. Hancock (1994). Growth and development of roots of grapevine (Vitis vinifera L.) in relation to water uptake from soil. In: Annals of Botany 74.1, pp. 75–85.

McCormack, M. L., E. Crisfield, B. Raczka, F. Schnekenburger, D. M. Eissenstat, and E. A. Smith-wick (2015). Sensitivity of four ecological models to adjustments in fine root turnover rate. In: Ecological modelling 297, pp. 107–117.

McDowell, N., W. T. Pockman, C. D. Allen, D. D. Breshears, N. Cobb, T. Kolb, J. Plaut, J. Sperry, A. West, D. G. Williams, et al. (2008). Mechanisms of plant survival and mortality during drought: why do some plants survive while others succumb to drought? In: New phytologist 178.4, pp. 719–739.

McDowell, N. G., M. Ball, B. Bond-Lamberty, M. L. Kirwan, K. W. Krauss, J. P. Megonigal, M. Mencuccini, N. D. Ward, M. N. Weintraub, and V. Bailey (2022). Processes and mechanisms of coastal woody-plant mortality. In: Global Change Biology 28.20, pp. 5881–5900.

Meunier, F., V. Couvreur, X. Draye, M. Zarebanadkouki, J. Vanderborght, and M. Javaux (2017). Water movement through plant roots–exact solutions of the water flow equation in roots with linear or exponential piecewise hydraulic properties. In: Hydrology and Earth system sciences 21.12, pp. 6519–6540.

Morandage, S., A. Schnepf, D. Leitner, M. Javaux, H. Vereecken, and J. Vanderborght (2022). Correction to: Parameter sensitivity analysis of a root system architecture model based on virtual field sampling. In: Plant and Soil 477.1, pp. 849–850. issn: 1573-5036. doi: 10.1007/s11104-021-05089-3. url: 10.1007/s11104-021-05089-3.

Morris, E. C., M. Griffiths, A. Golebiowska, S. Mairhofer, J. Burr-Hersey, T. Goh, D. Von Wangenheim, B. Atkinson, C. J. Sturrock, J. P. Lynch, et al. (2017). Shaping 3D root system architecture. In: Current biology 27.17, pp. 919–930.

Munnè-Bosch, S. (2014). Perennial roots to immortality. In: Plant Physiology 166.2, pp. 720–725.

Pierret, A., C. Doussan, Y. Capowiez, F. Bastardie, and L. Pagès (2007). Root functional architecture: a framework for modeling the interplay between roots and soil. In: Vadose Zone Journal 6.2, pp. 269–281.

Poppenwimer, T., I. Mayrose, and N. DeMalach (2023). Revising the global biogeography of annual and perennial plants. In: Nature 624.7990, pp. 109–114.

R Core Team (2024). R: A Language and Environment for Statistical Computing. R Foundation for Statistical Computing. Vienna, Austria. url: https://www.R-project.org/.

Roumet, C., C. Urcelay, and S. Díaz (2006). Suites of root traits differ between annual and perennial species growing in the field. In: New phytologist 170.2, pp. 357–368.

RStudio Team (2020). RStudio: Integrated Development Environment for R. RStudio, PBC. Boston, MA. url: http://www.rstudio.com/.

Saint Cast, C., C. Meredieu, P. Défossez, L. Pagès, and F. Danjon (June 2019a). Modelling root system development for anchorage of forest trees up to the mature stage, including acclimation to soil constraints: the case of Pinus pinaster. In: Plant and Soil 439. doi: 10.1007/s11104-019-04039-4.

Saint Cast, C., C. Meredieu, P. Dèfossez, L. Pagès, and F. Danjon (2019b). Modelling root system development for anchorage of forest trees up to the mature stage, including acclimation to soil constraints: the case of Pinus pinaster. In: Plant and Soil 439.1, pp. 405–430.

Schnepf, A., D. Leitner, M. Landl, G. Lobet, T. H. Mai, S. Morandage, C. Sheng, M. Zörner, J. Vanderborght, and H. Vereecken (2018). CRootBox: a structural–functional modelling framework for root systems. In: Annals of botany 121.5, pp. 1033–1053.

Scott, E. I., E. Toensmeier, F. Iutzi, N. A. Rosenberg, S. T. Lovell, N. R. Jordan, T. E. Peters, E. Akwii, and E. M. Broad Leib (2022). Policy pathways for perennial agriculture. In: Frontiers in Sustainable Food Systems 6, p. 983398.

Sevanto, S., N. G. Mcdowell, L. T. Dickman, R. Pangle, and W. T. Pockman (2014). How do trees die? A test of the hydraulic failure and carbon starvation hypotheses. In: Plant, cell & environment 37.1, pp. 153–161.

Steng, K., L. Fichtl, M. Athmann, A. Gattinger, and J. Döring (2026). Long-term vineyard management systems shape soil moisture dynamics, nitrogen cycling, and grapevine root distribution. In: European Journal of Agronomy 180, p. 128201. issn: 1161-0301. doi: 10.1016/j.eja.2026.128201. url: https://www.sciencedirect.com/science/article/pii/S1161030126002200.

Strock, C. F. and J. P. Lynch (2020). Root secondary growth: an unexplored component of soil resource acquisition. In: Annals of Botany 126.2, pp. 205–218.

Sun, K., M. Luke McCormack, L. Li, Z. Ma, and D. Guo (2016). Fast-cycling unit of root turnover in perennial herbaceous plants in a cold temperate ecosystem. In: Scientific Reports 6.1, p. 19698.

Therneau, T. M. (2026). A Package for Survival Analysis in R. R package version 3.8-6. url: https://CRAN.R-project.org/package=survival.

Tron, S., G. Bodner, F. Laio, L. Ridolfi, and D. Leitner (2015). Can diversity in root architecture explain plant water use efficiency? A modeling study. In: Ecological Modelling 312, pp. 200–210. doi: 10.1016/j.ecolmodel.2015.05.028.

Tumber-Dávila, S. J., H. J. Schenk, E. Du, and R. B. Jackson (2022). Plant sizes and shapes above and belowground and their interactions with climate. In: New Phytologist 235.3, pp. 1032–1056.

Vanderborght, J., V. Couvreur, M. Javaux, D. Leitner, A. Schnepf, and H. Vereecken (2024a). Mechanistically derived macroscopic root water uptake functions: The α and ω of root water uptake functions. In: Vadose Zone Journal 23.4, e20333.

Vanderborght, J., D. Leitner, A. Schnepf, V. Couvreur, H. Vereecken, and M. Javaux (2024b). Combining root and soil hydraulics in macroscopic representations of root water uptake. In: Vadose Zone Journal 23.3, e20273.

Vercambre, G., L. Pagès, C. Doussan, and R. Habib (2003). Architectural analysis and synthesis of the plum tree root system in an orchard using a quantitative modelling approach. In: Plant and Soil 251.1, pp. 1–11.

Waite, H., M. Whitelaw-Weckert, and P. Torley (2015). Grapevine propagation: principles and methods for the production of high-quality grapevine planting material. In: New Zealand Journal of Crop and Horticultural Science 43.2, pp. 144–161. doi: 10.1080/01140671.2014.978340. eprint: https://rsnz.onlinelibrary.wiley.com/doi/pdf/10.1080/01140671.2014.978340. url: https://rsnz.onlinelibrary.wiley.com/doi/abs/10.1080/01140671.2014.978340.

Weibull, W. (1939). A statistical theory of strength of materials. In: IVB-Handl.

Wessolek, G., M. Kaupenjohann, and M. Renger, eds. (2009). Bodenphysikalische Kennwerte und Berechnungsverfahren für die Praxis. German. Vol. 40. Bodenökologie und Bodengenese. 80 S. Berlin: Technische Universität Berlin, Selbstverlag.

Wickham, H. (2016). ggplot2: Elegant Graphics for Data Analysis. Springer-Verlag New York. isbn: 978-3-319-24277-4. url: https://ggplot2.tidyverse.org.

Zhou, X.-R., A. Schnepf, J. Vanderborght, D. Leitner, A. Lacointe, H. Vereecken, and G. Lobet (2020). CPlantBox, a whole-plant modelling framework for the simulation of water-and carbon-related processes. In: in silico Plants 2.1, diaa001.

